# Perinatal IL-1β-induced Inflammation Suppresses Tbr2+ Intermediate Progenitor Cell Proliferation in the Developing Hippocampus accompanied by Long-Term Behavioral Deficits

**DOI:** 10.1101/2020.05.08.084129

**Authors:** Stephanie Veerasammy, Juliette Van Steenwinckel, Tifenn Le Charpentier, Joon Ho Seo, Bobbi Fleiss, Pierre Gressens, Steven W. Levison

**Author notes:** **Corresponding Author:** Steven W. Levison, Ph.D., Director, Laboratory for Regenerative Neurobiology, Department of Pharmacology, Physiology, and Neuroscience, Rutgers New Jersey Medical School, 205 South Orange Ave., Newark, NJ 07103.

## Abstract

Meta-analyses have revealed associations between the incidence of maternal infections during pregnancy, premature birth, smaller brain volumes, and subsequent cognitive, motor and behavioral deficits as these children mature. Inflammation during pregnancy in rodents produces cognitive and behavioral deficits in the offspring that are similar to those reported in human studies. These deficits are accompanied by decreased neurogenesis and proliferation in the subgranular zone (SGZ) of the dentate gyrus (DG) of the hippocampus. As systemically administering interleukin-1 β (IL-1β) to neonatal mice recapitulates many of the brain abnormalities seen in premature babies including developmental delays, the goal of this study was to determine whether IL-1-mediated neuroinflammation would affect hippocampal growth during development to produce cognitive and behavioral abnormalities. For these studies, 10 ng/g IL-1β was administered twice daily to Swiss Webster mice during the first 5 days of life, which increased hippocampal levels of IL-1α and acutely reduced the proliferation of Tbr2^+^ neural progenitors in the DG. *In vitro*, both IL-1α and IL-1β produced G1/S cell cycle arrest that resulted in reduced progenitor cell proliferation within the transit amplifying progenitor cell cohort. By contrast, IL-1β treatment increased neural stem cell frequency. Upon terminating IL-1β treatment, the progenitor cell pool regained its proliferative capacity. An earlier study that used this *in vivo* model of perinatal inflammation showed that mice that received IL-1β as neonates displayed memory deficits which suggested abnormal hippocampal function. To evaluate whether other cognitive and behavioral traits associated with hippocampal function would also be altered, mice were tested in tasks designed to assess exploratory and anxiety behavior as well as working and spatial memory. Interestingly, mice that received IL-1β as neonates showed signs of anxiety in several behavioral assays during adolescence that were also evident in adulthood. Additionally, these mice did not display working memory deficits in adulthood, but they did display deficits in long-term spatial memory. Altogether, these data support the view that perinatal inflammation negatively affects the developing hippocampus producing behavioral deficits that persist into adulthood. These data provide a new perspective into the origin of the cognitive and behavioral impairments observed in prematurely-born sick infants.

**Highlights:**

- Systemic inflammation during the first 5 days of life increases hippocampal pro-inflammatory cytokines and chemokines
- Neuroinflammation reduces Tbr2^+^ neural progenitor cell proliferation in the dentate gyrus.
- IL-1 arrests the cell cycle of Tbr-2^+^ progenitors and enriches for neural stem cells.
- Adolescent mice that experienced neonatal systemic inflammation have a persistent anxiety disorder.
- Mice that experienced neonatal systemic inflammation have deficits in spatial memory as adults.

## Introduction

Maternal fever or other evidence that a pregnant woman has an underlying infection is strongly associated with premature birth and a poor neurological assessment of the newborn, often predicting future disorders such as attention hyperactivity disorder (ADHD), autism, learning disabilities and cerebral palsy (Hagberg *et al.*, 2015). Meta-analyses have revealed associations between the incidence of premature birth (birth at <37 of 40 weeks gestation), smaller total brain volumes, and cognitive, motor, and behavioral deficits in young and adolescent children (Aarnoudse-Moens *et al.*, 2009; Bhutta *et al.*, 2002; de Kieviet *et al.*, 2012). Therefore, the high incidence of infection in premature infants has warranted additional studies into the causal relationship between infection and the constellation of grey and white matter neuropathologies observed in preterm born infants termed Encephalopathy of Prematurity (EoP). To date, most basic science and clinical research has focused on structural changes to the neocortex and periventricular white matter as the cause of disabilities following EoP. We have developed a complementary premise, which is that infection in the sick preterm infant compromises the development and function of the hippocampus, a structure long known to be essential for human learning, memory and stress responses. This hypothesis is supported by neuroimaging studies which show that babies born very prematurely have smaller and abnormally-shaped hippocampi at term-equivalent age compared to babies born at term suggesting that the development of the hippocampus is delayed. These same children had deficits in memory tasks even after adjusting for relevant perinatal, sociodemographic, and developmental factors when tested at 7 and in a separate study at 19-20 years of age (Aanes *et al.;* Beauchamp *et al.*, 2008).

The hippocampus is a bilateral brain structure that is recognized as essential for learning, memory, spatial navigation, and responses to stressful stimuli. While the pyramidal cell layer of the Cornu Ammonis of term infants is cytoarchitecturally similar to that seen in the adult, the granule cell layer of the DG of the newborn infant is immature and it continues to grow after birth, increasing by ~50% during the 1^st^ 3 months of post-natal life (Arnold and Trojanowski, 1996; Holland *et al.*, 2014). Studies of the neural progenitors (NPs) of the DG suggest that these cells are bipotential, giving rise to both neurons and astrocytes. NPs of the developing hippocampus include quiescent neural progenitors (Type 1) and transit-amplifying progenitors (TAPs). The TAPs comprise amplifying neural progenitors which are non-fate-determined (Type 2a) and fate-determined (Type 2b). TAPs then give rise to neuroblasts (Type 3), which are neuronally restricted, but capable of mitosis. Once neuroblasts lose their ability to divide, they become immature neurons that fully differentiate into mature, glutamatergic granule cell neurons (Hsieh, 2012); it takes approximately 30 days for a mature neuron to be produced from a Type 2a NP (Encinas *et al.*, 2011). After producing multiple generations of neurons, the Type 1 and Type 2a progenitors generate astrocytes.

Decreased hippocampal neurogenesis has been reported in murine models of prenatal inflammation using lipopolysaccharide (LPS) or polyinosinic:polycytidylic acid (Poly I:C) to model inflammation caused by infection with gram-negative bacteria or viruses, respectively (Cui *et al.*, 2009; Hofer *et al.*, 2011; Meyer *et al.*, 2006; Smith *et al.*, 2014; Wolf *et al.*, 2011). The decreased neurogenesis seen in these models may be due to reduced proliferation of the neural stem cells (NSCs) or TAPs and a subsequent reduction in differentiated neurons; however, this has not been established. Interestingly, rats and mice exposed to LPS and poly I:C also show cognitive and behavioral deficits at later stages, similar to those reported in clinical studies of preterm infants exposed to inflammatory insults.

Multiple cytokines and chemokines, including IL-1β, IL-6, and TNFα are produced with *in utero* infection in humans and rodents (Allred *et al.*, 1997; Arrode-Bruses and Bruses, 2012; Malaeb and Dammann, 2009). IL-1 includes two cytokines encoded by separate genes; interleukin-1α (IL-1α) and interleukin-1β (IL-1β). IL-1α is produced as a 31 kDa pro-peptide that is enzymatically cleaved to a 17kDa biologically active form that binds to IL-1R to activate cells. Similarly, IL-1β is expressed in the cytoplasm as a 31 kDa pro-form and is processed to a 17 kDa peptide involving IL-1β converting enzyme, also known as caspase-1. In rodent studies, both maternal and neonatal injections of LPS and poly I:C elevate levels of IL-1 in the pups, which in turn correlates with abnormal brain development (Arrode-Bruses and Bruses, 2012; Bilbo *et al.*, 2018; Boksa, 2010). One reason that the hippocampus is especially sensitive to high levels of IL-1 is that the IL-1 receptor is expressed at extraordinarily high levels within the DG of the hippocampus (Cunningham *et al.*, 1992). The regulated expression of both IL-1R1 (Simao *et al.*, 2016) and IL-1β during early postnatal development of the hippocampus (Garay *et al.*, 2013) suggests that this cytokine may have an important role during hippocampal development.

*In vitro* studies have shown that IL-1β can negatively affect hippocampal neurogenesis. Koo and Duman 2008 performed *in vitro* studies of adult rat hippocampal NPs and showed that NPs express IL-1R1 and that activation of IL-1R1 by IL-1β decreased NP proliferation via NF-κB signaling (Koo and Duman, 2008). Similarly, *in vitro* studies on embryonic rat hippocampal NPs by Green and colleagues revealed decreased NP proliferation and decreased neuronal differentiation as a result of IL-1β signaling through IL-1R1 (Green *et al.*, 2012), supporting the conclusion that IL-1R1 signaling negatively affects hippocampal NPs. However, in both of these studies, supraphysiological concentrations of IL-1β were used; therefore, their results are suspect. Furthermore, McPherson and colleagues performed parallel studies in adolescent vs. adult mice after a chemically induced injury to the hippocampus. They established that IL-1R1 signaling was prominent in adolescents whereas IL-6 signaling was predominant in adult mice (McPherson *et al.*, 2011). Contrary to other studies, these *in vitro* results showed increased proliferation of NPs after treatment with IL-1α, which questions the role of IL-1R1 signaling on the proliferation of immature hippocampal NPs. Collectively, these studies warrant further investigation into the role of IL-1R1 ligands in hippocampal neurogenesis and how this interaction may contribute to cognitive, motor, and behavioral deficits seen in children and adolescents born prematurely (Aarnoudse-Moens *et al.*, 2009; Bhutta *et al.*, 2002; de Kieviet *et al.*, 2012).

Mice are born precociously, and though it is difficult to compare the development of the lissencephic brain of the mouse to the gyrencephalic human brain, comparative studies that have evaluated multiple criteria that include birthdates of specific neuronal subpopulations, involution of the ventricular zone, onset of myelination, neurochemical levels, metabolism and EEG patterns, conclude that the postnatal ages of P1 to P5 in the mouse represent the period from gestational week 23 to gestational day 32 in the human (Semple *et al.*, 2013). Therefore, for these studies where our goal was to model EOP and to evaluate the effects on hippocampal development, we chose to use the EOP paradigm established by Favrais et al., (2011) where recombinant IL-1β is administered twice daily from postnatal day 1 (P1) to P4 and once on P5.

## Materials and Methods

### Animals and Drug Administration

All experiments were performed in accordance with the research guidelines set forth by the Institutional Animal Care and Use Committee (IACUC) of Rutgers New Jersey Medical School and were in accordance with National Institute of Health Guide for the Care and Use of Laboratory Animals (NIH publication No. 80-23) revised in 1996 and the Animal Research: Reporting of In Vivo Experiments (ARRIVE) guidelines. Sample sizes per experiment were chosen to achieve sufficient statistical power with minimal numbers of animals based on pilot studies. Every experiment used pups from more than two litters with each treatment group comprised of mice derived from 2 to 6 litters. When only 3 pups were analyzed, each pup was from a different litter. Timed pregnant Swiss Webster and C57Bl/6 mice (Charles River Laboratories, Wilmington, MA) were group housed and kept on a 12-hour light:dark cycle (lights turned on at 7:00 am and turned off at 7:00 pm) with ad libitum access to food and autoclaved water. Sex was determined at P2 and confirmed by abdominal examination at death. Both male and female mouse C57BL/6 or Swiss Webster pups were used for *in vitro* studies. Only male Swiss Webster pups were used for histological and behavioral studies, as females have been previously shown to be much less affected using this model of perinatal systemic inflammation (Favrais *et al.*, 2011). On the day after birth, the litters were culled to contain only male pups. Animals were randomly assigned to either the IL-1ß or the PBS group. Mice were administered either a 5 μL volume of phosphate-buffered saline (PBS) containing 10 μg/kg of carrier-free rmIL-1β (R&D Systems, Minneapolis, MN) or PBS alone (control) twice a day via the intraperitoneal (IP) route on days P1 to P4 and once on day P5 (the day of birth was defined as P0). The mice received Pelikan black ink tatoos into their left (IL-1ß) or right (PBS) footpads for identification purposes.

### RNA extraction and quantification of gene expression by real-time qPCR

Total RNA from mouse hippocampi was extracted with the RNeasy mini kit according to the manufacturer’s instructions (Qiagen, Courtaboeuf, France). RNA quality and concentration were assessed by spectrophotometry with the NanodropTM apparatus (Thermoscientific, Wilmington, DE, USA). Total RNA (1 μg) was reverse transcribed using the iScriptTM cDNA synthesis kit (Bio-Rad). RT-qPCR was performed in duplicate for each sample using SYBR Green Super-mix (Bio-Rad) for 40 cycles with a 2-step program (5 s of denaturation at 96 °C and 10 s of annealing at 60 °C). Amplification specificity was assessed with a melting curve analysis. Primers were designed using Primer3 software. The relative expression of genes of interest were expressed relative to expression of the reference gene, Glyceraldehyde 3-phosphate dehydrogenase (GAPDH), which an earlier study found was unaffected in this model (Van Steenwinckel *et al.*, 2019). Analyses were performed with the Biorad CFX manager 2.1 software.

### Neurosphere Cultures

Neonatal C57BL/6 or Swiss Webster mouse pups at P5 were euthanized by decapitation, and their brains were immediately removed. The brains were cut along the longitudinal fissure and the hippocampus was dissected out of each hemisphere. The tissue was enzymatically digested in DMEM/F-12 medium containing 0.1% trypsin (Sigma-Aldrich), 10 units/mL papain (Worthington Biochemical Corporation, Lakewood, NJ), 40 μg/mL DNase I (Sigma-Aldrich), and 3 mM MgSO4 (Sigma-Aldrich) in DMEM/F-12 for 5 min at 37°C with intermittent shaking. The enzymatic reaction was stopped using an equivalent volume of 10% newborn calf serum Hyclone (Logan, UT) in DMEM/F-12. The tissue was centrifuged at 800 rpm for 5 min followed by a wash with DMEM/F-12 at 1200 rpm for 5 min, after which the pellet was resuspended in 3 mL of DMEM/F-12, mechanically triturated until smooth. The cell suspension was then filtered through a 40 μm cell sieve (BD Falcon, Franklin Lakes, NJ) and pelleted at 1200 rpm for 5 min. The pellet was then resuspended in neurosphere growth medium, and cells were plated at a density of 1.75e5 cells/mL with 200 μL/cm^2^ in either 60 mm or 100 mm tissue culture dishes. The neurosphere growth medium consisted of DMEM/F-12 supplemented with GlutaMax, gentamycin, B27 supplement (Invitrogen, Waltham, MA), 20 ng/mL recombinant human epidermal growth factor (EGF, PeproTech, Rocky Hill, NJ) and 20 ng/mL recombinant human fibroblast growth factor-2 (FGF-2, Alomone Labs, Jerusalem, Israel). Plates were incubated at 37°C with humidified 5% CO2 to permit primary neurosphere formation and were fed every 2 days by replacing half of the medium with an equal volume of fresh growth medium containing 2X EGF and FGF-2. Primary neurospheres were collected after 7 days by centrifugation at 1200 rpm for 5 min. They were dissociated for 5 min at 37°C in 70% Accutase (EMD Millipore, Billerica, MA) in DMEM/F-12 and plated into either 60 mm or 100 mm tissue culture dishes as described above with the addition of rmIL-1α or rmIL-1β (50 or 500 pg/mL, R&D Systems, Minneapolis, MN)) for treatment groups. Cells were fed every 2 days by replacing half of the medium with an equal volume of fresh growth medium containing 2X EGF and FGF-2 and additional IL-1 in treated plates. Secondary neurosphere number and size were assessed after 6 days of growth using phase contrast images taken at 4X magnification. Spheres were dissociated with 70% Accutase as described above to enable quantitation of total cell yield from each treatment group. Neurosphere size was measured using Zeiss Axiovision (Carl Zeiss, Oberkochen, Germany) software. Spheres with diameters ≥ 30 μm were counted as neurospheres. Results of the neurosphere counts, size, and cell yield were expressed as mean ± SEM and statistical analyses were performed using unpaired t-test or ANOVA followed by Bonferroni’s multiple comparisons post-hoc test in GraphPad Prism (GraphPad Software Inc., La Jolla, CA).

### Self-Renewal Studies

Primary neurospheres were grown in neurosphere growth medium in the presence versus absence of IL-1β (500 pg/mL) (treated versus control, respectively) in 100 mm tissue culture dishes. The medium was changed and supplemented with IL-1β every 2 days, as described above. After 7 days of growth, neurospheres were collected, dissociated with Accutase and plated at a density of 1.75e^5^ cells/mL of neurosphere growth medium with 200 μL/cm^2^ in 60 mm tissue culture dishes. Neurospheres were passaged every 7 days as described and total cell yield was recorded at each passage.

### Limiting Dilution Analysis

Primary neurospheres were grown in neurosphere growth medium in the presence versus absence of IL-1β (500 pg/mL) (treated versus control, respectively) in 100 mm tissue culture dishes. The medium was changed and supplemented with IL-1β every 2 days, as described above. After 7 days of growth, neurospheres were collected and dissociated with 70% Accutase and plated at reducing densities in 96-well plates with 200 μL of growth medium. Cells were plated at the following densities with 8 replicate wells per density: 1000, 500, 250, 188, 125, 94, 63, 47, 31, and 16 cells/well. Half of the volume of media in each well was replaced with fresh growth medium containing 2X EGF and FGF-2 every 2 days. After 9 days of growth, the fraction of wells negative for neurospheres was quantified. These data were then log transformed and plotted against plating density. A linear regression was performed on the data using GraphPad Prism.

### Cell Cycle Analyses

Primary neurospheres were grown for 4 days. The medium was replaced with growth factor-free medium for 15 hours to synchronize the cells in the cell cycle. The neurosphere growth medium containing growth factors was then provided in the presence versus absence of IL-1α or IL-1β (500 pg/mL). Additions of 500 pg/mL IL-1 were added to plates every 12 h, and neurospheres were collected, dissociated on ice, and fixed with 1% PFA at 12, 18, 24 and 48 h post-recovery/ treatment. Cells were incubated on ice in rabbit polyclonal anti-Tbr2 antibodies for 30 min, washed 3X with buffer, followed by Alexa Fluor 700-conjugated goat anti-rabbit IgG (Molecular Probes, Carlsbad, CA, diluted at 1:1000) for 30 min followed by wash 3X with buffer. All staining and washes with antibodies were performed with 1X saponin/1% BSA in PBS (without Ca^2+^ or Mg^2+^) on ice or at 4°C. Cells were washed once with PBS, incubated on ice in 1 μg/mL 4’, 6’-diamidino-2-phenylindole (DAPI, Sigma Aldrich) in PBS containing 0.1% Triton-X-100 for 30 min, and then subjected to DNA content analysis on an LSRII flow cytometer (BD Biosciences, Franklin Lakes, NJ). A minimum of 10,000 cells was analyzed at each time point. Appropriate gating strategies were applied to exclude cell debris and aggregates. Flow cytometry data were analyzed using BD FACS Diva™ (BD Biosciences, Franklin Lakes, NJ) or ModFit™ software (Verity Software House, Topsham, ME).

### Western Blotting for Cell Cycle Proteins

Secondary neurospheres were grown for 4 days and then the medium was replaced with growth factor-free medium for 6 h to synchronize the cells in the cell cycle. Neurosphere growth medium containing growth factors was then provided in the presence versus absence of IL-1α or IL-1β (500 pg/mL). Neurospheres were collected in RIPA lysis buffer containing Phosphatase Inhibitor Cocktail 2 (Sigma-Aldrich) and cOmplete Mini Protease Inhibitor Cocktail (Roche Molecular Systems, Indianapolis, IN) at 0.5, 3, 6, 12, and 18 h post-recovery/ treatment and lysed on ice for 10 min. Lysates were stored at −20° C. Protein concentration was determined using the BCA assay and 30 μg protein per aliquot was stored at −80° C. Proteins were separated using 12% Bis-Tris Acetate gels (Invitrogen, Carlsbad, CA) with MOPS buffer (Invitrogen, Carlsbad, CA) and then transferred to 0.2 μm pore nitrocellulose membranes (Invitrogen). The blots were probed with the following primary antibodies: rabbit antip21 (Santa Cruz Biotechnology Inc., Dallas, TX, catalog # SC-756, diluted at 1:200), mouse anti-p27 (BD Transduction Laboratories, San Jose, CA, catalog # 610242, diluted at 1:2500), rabbit anti-p57 (Santa Cruz Biotechnology Inc., Dallas, TX, catalog # SC-1040, diluted at 1:100), rabbit anti-p57 (Santa Cruz Biotechnology Inc., Dallas, TX, catalog # SC-6243, diluted at 1:200), mouse anti-Cyclin D1 (Santa Cruz Biotechnology Inc., Dallas, TX, catalog # SC-8396, diluted at 1:100), mouse anti-Cyclin B1 (Santa Cruz Biotechnology Inc., Dallas, TX, diluted at 1:2000), mouse anti-Cyclin B (BD Transduction Laboratories, San Jose, CA, diluted at 1:1000). The blots were also probed with primary antibodies against beta-actin (Sigma-Aldrich, St. Louis, MO, diluted at 1:100), which served as a loading control. HRP-conjugated goat anti-rabbit IgG (Jackson ImmunoResearch, diluted at 1:2500) and HRP-conjugated goat anti-mouse IgG (Jackson ImmunoResearch, diluted at 1:2500) were used to detect the primary antibodies. The Western Lightning ECL Pro kit (Perkin Elmer, Waltham MA) was used to visualize protein bands. Protein bands were detected and quantified using a Bio-Rad ChemiDoc Imaging System (Hercules, CA) and Lab Pro software (Vernier Software & Technology, Beaverton, OR).

### Multiplex Immunoassay

Pups received an overdose of pentobarbital and were then perfused with RPMI culture medium. Hippocampi were dissected at P2, P5, and P10 in control and IL-1β-treated groups. Samples were prepared and analyzed using a Bio-Plex Pro Mouse Cytokine 23-Plex Assay kit (Bio-Rad, Hercules, CA). Assays were performed according to the manufacturer’s instructions. Reagents were kept on ice until use, with minimal exposure of the beads to light. Corresponding buffer blanks were run to determine the level of background. Sensitivity of the assay was > 2.5 pg/mL for IL-1α, IL-6, VEGF-A and KC; greater than 13 pg/mL for MCP-1 and greater than 19 pg/mL for IL-1β. All samples were run in duplicate at 900 μg protein/well. Corresponding buffer blanks were run to determine the level of background. All the wash steps were performed on a Bio-Plex Pro Wash Station at room temperature. The plates were then read in the Bio-Plex 200 System and the data analyzed using Bio-Plex Manager 4.1 software.

### Immunofluorescence

For histology studies with paraffin-embedded samples, mice were deeply anesthetized with xylazine/ ketamine (10 mg/ 75 mg/kg) before intracardiac perfusion on P10, and the brains were removed and post-fixed in 4% paraformaldehyde in PBS, pH 7.4 for 4 days at room temperature. The brains were cut along the longitudinal fissure and then dehydrated through a series of alcohols and embedded in paraffin. The left hemispheres were sectioned sagitally at 8 μm with a microtome, mounted on Superfrost Plus slides (VWR, Radnor, PA), and stored at room temperature until staining. Three section pairs containing dorsal hippocampus were selected. Sections were encircled with barrier pen, blocked in TDB superblock (TBS containing 10% normal donkey serum and 0.1g/mL BSA) for 1 h followed by incubation overnight at 4°C in a humidified chamber in rabbit anti-Ki67 antibodies (eBioscience, Waltham, MA, catalog # 14-5698-82, diluted at 1:200) (diluted in 20% TDB superblock and 0.3% Triton-X100). Slides were washed 4X for 5 min each in T-Buffer (TBS containing 0.05% Triton-X100) followed by incubation in Alexa Fluor 647-conjugated donkey anti-rabbit IgG (Jackson ImmunoResearch, West Grove, PA, diluted at 1:200) for 2 h at room temperature in a humidified chamber. Slides were washed 4X for 5 min each in T-Buffer followed by a rinse in TBS for 5 min and then incubated in 1:5000 DAPI (Sigma) for 15 min at room temperature. Slides were washed 2X for 5 min in TBS, briefly rinsed in ddH2O, and mounted with coverslips using Fluoro-Gel. Images were collected on an Olympus BX51 microscope (Center Valley, PA) and captured by a Q-imaging Retiga-2000R CCD camera (Surrey, BC, Canada). Images of the hippocampi were captured using a 40X objective and were used for stereological counts of Ki67 in the SGZ using the physical dissector (Microbrightfield, Williston, VT).

For histology studies with cryopreserved samples, mice were deeply anesthetized with xylazine/ ketamine (10 mg/ 75 mg/kg) before intracardiac perfusion on P10 and P60, and brains were removed and drop-fixed in 4% PFA overnight at 4° C followed by cryoprotection in 30% sucrose for 2 days. Brains were briefly rinsed in PBS(no Ca^2+^ or Mg^2+^), blocked using a steel matrix and embedded in OCT (Tissue-Tek, München, Germany) using a dry ice/ ethanol bath. Whole brains were sectioned coronally at 40 μm with a cryostat, thaw-mounted on Superfrost Plus slides and stored at −20°C until staining. Three sections containing dorsal hippocampus were selected for the P10 study; six sections spanning the dorsal and ventral hippocampus were selected for the P60 study. Sections were encircled with a barrier pen, blocked in TDB superblock for 1 h followed by incubation in rabbit polyclonal anti-Tbr2 (Abcam, Cambridge, MA, catalog # ab23354, diluted at 1:500) or rabbit monoclonal anti-Tbr2 (Abcam, Cambridge, MA, catalog # ab183991, diluted at 1:1000), and rat anti-Ki67 (Vector Laboratories, Burlingame, CA, catalog # Z0311, diluted at 1:1000) at 4°C in a humidified chamber overnight. Slides were washed 3X for 10 min each in T-Buffer followed by one wash in TBS and incubation in Alexa Fluor 647-conjugated donkey anti-rabbit IgG (Jackson ImmunoResearch, diluted at 1:250) and Alexa Fluor 488-conjugated donkey anti-rat IgG (Jackson ImmunoResearch, diluted at 1:300) for 2 h at 37°C followed by 1 h at room temperature in a humidified chamber. Slides were washed 3X for 10 min each in T-Buffer followed by one wash in TBS for 5 min and then incubated in 1:5000 DAPI for 15 min at room temperature. Slides were washed 2X for 5 min in TBS, briefly rinsed in ddH2O, and mounted with coverslips using Fluoro-Gel (Electron Microscopy Services, Hatfield, PA). Image stacks of the SGZ were obtained on an Olympus BX51 microscope and captured by a Q-imaging Retiga-2000R CCD camera at 100X using an oil immersion objective. The optical fractionator was used to obtain stereological counts of Tbr2- and Ki67-positive cells. Fifty percent of the SGZ was counted for P10 animals. Live counting at 40X was used to perform optical fractionator stereological counts of Tbr2^+^ and Ki67^+^ cells in 100% of the SGZ for P60 animals.

All of the primary antibodies used were evaluated by Western blot and found to produce strong bands at the molecular weights predicted for the protein they were raised against. We found it necessary to evaluate several different anti-Tbr-2 antibodies before we found ones that gave consistently robust staining within the dentate gyrus. All secondary antibody combinations were carefully examined to ensure that there was no bleed through between fluorescent dyes or cross-reactivity between secondary antibodies. No signal above background was obtained when the primary antibodies were replaced with pre-immune sera. The Annexin V Apoptosis Detection Kit with PI was purchased from Biolegend (San Diego, CA).

### Elevated Plus Maze

Mice were subjected to a battery of behavioral test by an investigator who was blinded to the experimental groups. The elevated plus maze (EPM) was used to assess anxiety. The EPM apparatus consisted of 4 arms (two open without walls and two enclosed by 15.25 cm high walls) 30 cm long and 5 cm wide and elevated 90 cm off the ground. Open arms of the EPM were situated toward the wall and the observer. The protocol described by Walf et al., 2007 was followed for these studies (Walf and Frye, 2007). On the test day, mice at P36 were transferred to the testing room in their home cages and habituated for 30 minutes. Prior to testing each mouse, the base and walls of the EPM apparatus were wiped down with Labsan C-DOX disinfectant and allowed to dry thoroughly. Each mouse was gently picked up and carried by the investigator to the apparatus where they were placed in the center of the apparatus with the head oriented toward the open arm facing the wall. Each mouse was allowed to explore the arms of the apparatus for 5 minutes. Movements throughout the apparatus (identified as “center”, “closed arms”, and “open arms”) were recorded with an aerial camera using the ANY-maze behavioral tracking software (ANY-maze, Wood Dale, IL). After the test, the mouse was returned to its home cage.

### Open Field Task

The open field task tests for general locomotor activity and may also be used to assess anxiety. Locomotor activity was measured using the Photobeam Activity System (PAS)-Open Field apparatus (San Diego Instruments, San Diego, CA). On the test day, mice at P39 were transferred to testing room in their home cages and habituated for 30 minutes. Prior to testing each mouse, the glass base and clear acrylic walls of the chambers were wiped down with 70% ethanol and water, respectively, and allowed to dry thoroughly. Each mouse was gently picked up and carried by the investigator to the apparatus where it was placed into the center of the chamber. Locomotor activity was measured across 12 consecutive intervals of 5 minutes each, for a total of 60 minutes of testing with the investigator absent from the testing room. After the test, each mouse was returned to its home cage. Parameters were measured by two stacked sets of 16 x 16 photobeams and the PAS-Open Field software, and included distance travelled, average speed, total resting time, rearing, time spent in center (defined as area not within 4 photobeams from the edges of the chamber) and number of entries into the center.

### T-Maze Forced Alternation

The T-maze forced alternation task is used to test working memory. The task employs a mouse T-maze apparatus with a stem length of 35 cm, arm lengths of 28 cm each, lane width of 5 cm and wall height of 10 cm (ANY-maze, Wood Dale, IL). The protocol described by Lalonde et al. 2002 was followed (Lalonde, 2002). On the test day, mice at P60 were transferred to the testing room in their home cages and habituated for 30 minutes. Prior to testing each mouse, the metal base and opaque acrylic walls of the apparatus were wiped with Labsan C-DOX disinfectant solution and allowed to dry thoroughly. In the forced choice phase, the entrance of one arm of the apparatus was blocked. Each mouse was gently picked up and carried by the investigator to the apparatus where they were placed at the starting point of the T-maze stem. Once the mouse entered the open arm of the maze, the arm entrance was blocked, restricting the mouse in the arm. After 45 sec, the mouse was gently removed from the arm and sequestered away from the apparatus for 60 sec before being reintroduced to the maze for the preferred choice phase. In the preferred choice phase, the entrances of both arms of the maze were left open and the mouse was once again placed at the starting point of the T-maze stem. The mouse was allowed to explore the maze and the arm entered was recorded. After each test, each mouse was returned to the home cage. If the mouse entered the arm that was blocked during the forced choice phase, the trial was recorded as an “alternation”. If the mouse entered the arm that was open during the forced choice phase, the trial was recorded as a “failure”. Each mouse was tested across four trials in one day with an inter-trial interval of 1 h. To ensure that the preferred choice was not learned across the trials, the arm that was blocked during the forced choice phase was changed for each trial. The percent alternation was calculated for each mouse across all trials.

### Barnes Maze

The Barnes maze task tests spatial learning and memory and requires a total of 6 testing days (Days 1 through 4 are training days and Days 5 and 12 are probe trial days). Mice began training on P65 and completed the second probe trial on P76. The task employed a 90 cm diameter circular acrylic platform painted black with 16-5 cm diameter holes equally spaced 2.5 cm from the periphery of circular edge of the platform. The platform was secured on top of a portable barstool 76 cm from the floor and placed in the center of a room with high contrast extra-maze spatial cues of varying shapes and patterns scattered throughout a brightly lit room. One of the holes was equipped with a small escape box containing fresh aspen bedding for each trial. The escape hole, box, and the orientation of the maze in the room were kept consistent throughout the training days. The protocol described by Patil et al 2009 was followed (Patil *et al.*, 2009). One each test day, mice were transferred to the testing room in their home cages and habituated for 30 minutes in the far corner of the room away from the apparatus and cues. Prior to testing each mouse, the platform of the apparatus was wiped with Labsan C-DOX disinfectant solution and allowed to dry thoroughly. On Days 1 through 4, each mouse was placed in the center of the platform in a transport box for 10 seconds. The transport box was then removed, and the mouse was allowed to explore the platform until it entered the escape hole target. The mice were kept in the escape box for 60 sec and then gently removed and returned to their home cage. Each mouse was trained across 4 trials per day with an inter-trial interval of 12 minutes. On Days 5 and 12, the escape box was removed from the apparatus and the maze was rotated to avoid use of intra-maze cues. Each mouse was introduced to the apparatus as described above for one trial; however, they were allowed to explore the platform for 90 sec after which they were gently removed from the platform and returned to their home cage. Movements throughout the platform were recorded with an aerial camera using the ANY-maze behavioral tracking software (ANY-maze, Wood Dale, IL). The average number of errors, defined as head pokes into any hole other than the target, per trial were recorded for each training day per mouse. The hole identity and number of errors were recorded for each mouse during each probe trial.

### Data Analysis and Statistics

Data were imported into Prism 8 (GraphPad Software, La Jolla, CA, USA) for statistical analyses using unpaired t-test, one-way ANOVA followed by Tukey’s multiple comparisons test, or two-way ANOVA followed by either Tukey’s or Sidak’s multiple comparisons test. For all statistical tests, the significance level was set at α=0.05. Graphs were produced in Prism 8, BD FACS Diva, or ModFit, and error bars denote standard error of the mean (SEMs).

### Data Availability

All new data are available from the authors upon request.

## Results

### Pro-inflammatory cytokines and chemokines are elevated in the hippocampus of mice after neonatal systemic inflammation

To determine whether hippocampal cytokine expression was altered in mice experiencing neonatal systemic inflammation during the first 5 days of life, cohorts of mice at P2, P5, and P10 were euthanized and protein levels of FGF-2, IFN-γ, IL-1α, IL-1β, IL-6, IL-10, IL-12p40, IL-17, IL-18, KC/CXCL1, MCP-1/CCL2, MIP-1β/CCL4, TNF-α, and VEGF-A were determined using a cytokine multiplex immunoassay on whole hippocampal lysates. Increases in MCP-1/CCL2 at P2 (q(20)=8.252, p<0.001, Figure 1B) and P5 (q(20)=4.876, p<0.05, Figure 1B), KC/CXCL1 at P2 (q(20)=5.473, p<0.05, Figure 1C), and VEGF-A at P5 (q(20)=5.446, p<0.05, Figure 1D) were detected by a two-way ANOVA followed by a Tukey’s multiple comparisons test comparing treatment groups and timepoints. Systemic IL-1β increased IL-1α (t(20)=2.635, p<0.05, Figure 1A) and VEGF-A (t(20)=2.822, p<0.05, Figure 1D) levels compared to PBS treated animals at P10 as detected by a Sidak’s multiple comparisons test comparing treatment groups at each individual timepoint. Developmental increases in the levels of IL-1a and IL-6 also were observed. No change in levels of the other cytokines or of IL-6 or IL-1β were detected between control and IL-1β-treated pups (Figure 1E,F).

**Figure 1.**
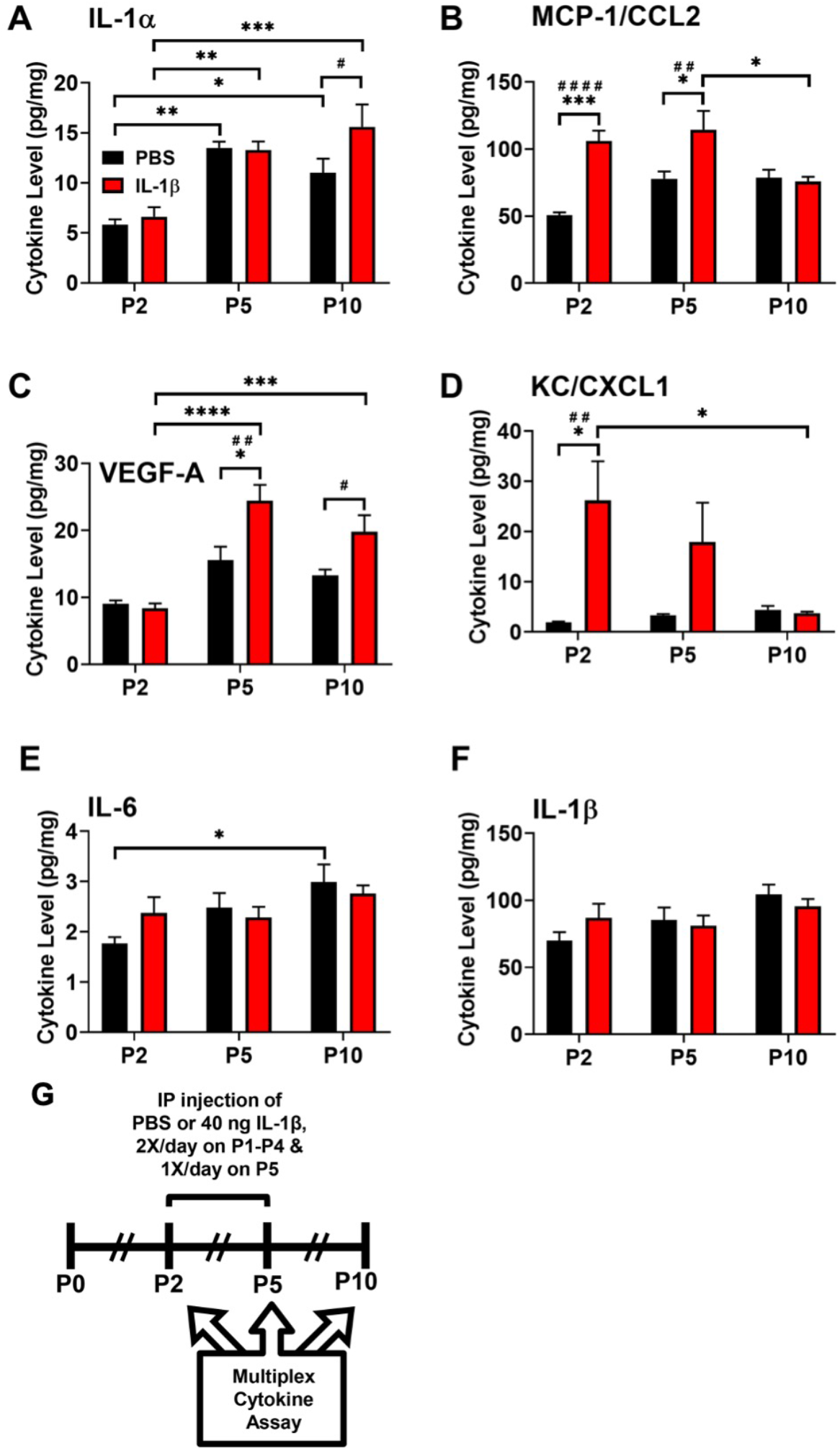
IL-1α, CCL2, VEGF-A and CXCL1 protein levels are significantly elevated acutely and subacutely in the hippocampus of mice with systemic inflammation. Neonatal male Swiss Webster mice were administered PBS or IL-1β from P1 through P5 and euthanized at P2, P5, and P10 after perfusing the animals with heparinized RPMI culture medium. Whole hippocampal tissue lysates were assayed for cytokine protein levels of (A) IL-1α, (B) MCP-1/CCL2 (C) VEGF-A. and (D) KC/CXCL1, (E) IL-6 and (F) IL-1β. Results are expressed as mean pg/mg total protein ± SEM; *p<0.05, ** p< 0.01, ***p<0.001, ****p<0.0001 by two-way ANOVA with Tukey’s multiple comparisons (*s)and Sidak’s tests (#s), n=5 mice per group at P2, n=4 mice per group at P5 and P10. (G) Timeline of treatment and endpoints for *in vivo* multiplex cytokine assay.

### The proliferation of TAPs is acutely reduced in mice with systemic inflammation

An earlier studied showed that male mice receiving systemic administration of IL-1β from P1 through P5 displayed cognitive deficits in two memory tasks (Favrais *et al.*, 2011). To determine if these memory deficits could be due to changes in the development of the DG of the hippocampus, a region of rapid growth in neonates, mice were administered IL-1β from P1 through P5 and euthanized at P10 (Figure 2A). Sections were stained for Ki67 as a marker for cycling cells in the SGZ (Figure 2B-C) and the physical dissector was used to count the number of cycling cells. A 35% decrease in the number of Ki67^+^ cells was found in the dorsal hippocampi of mice receiving IL-1β versus PBS, as determined by an unpaired t-test (t(4)=3.258, p=0.031, Figure 2D).

**Figure 2.**
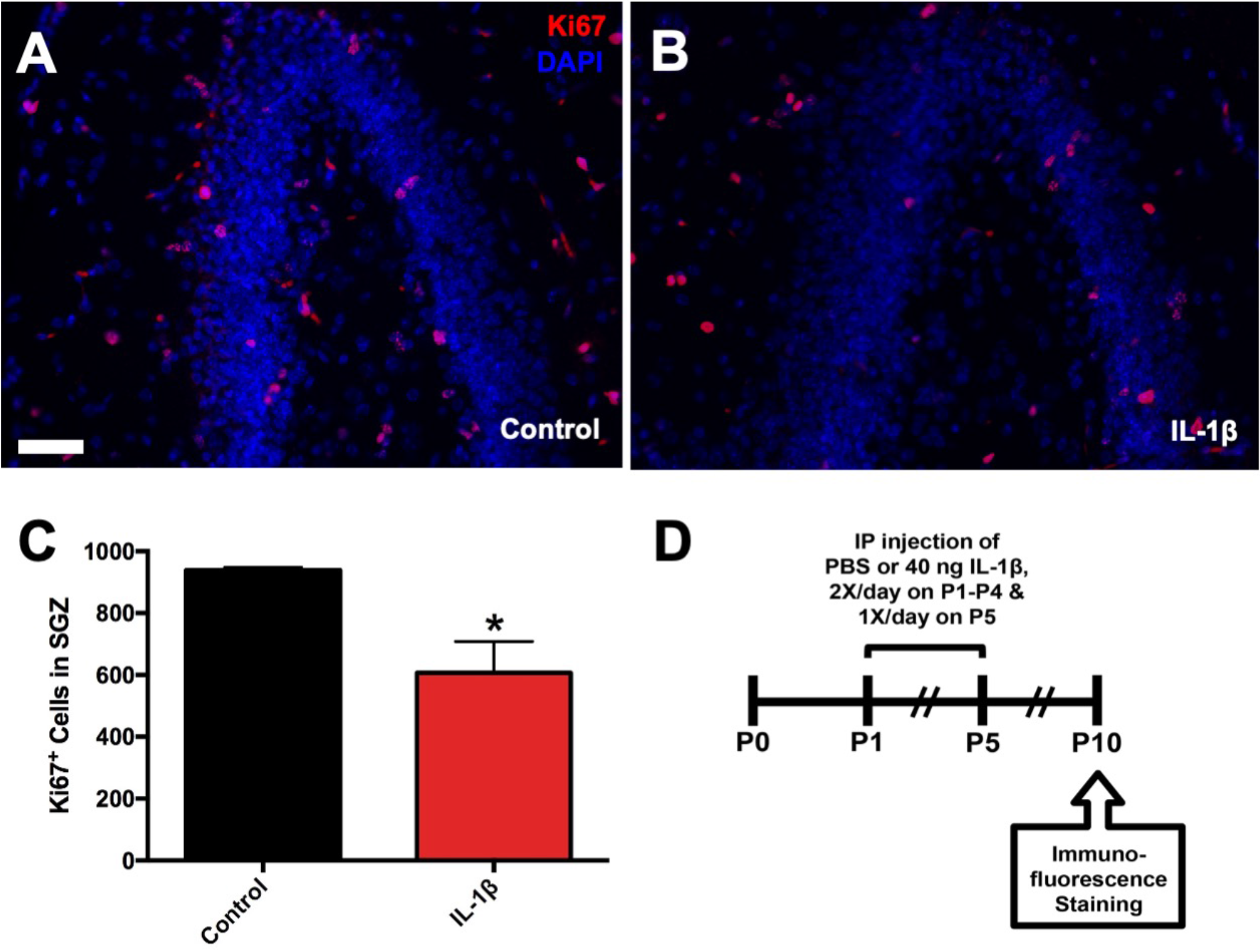
The number of cycling cells in the subgranular zone is reduced in mice with systemic inflammation. Neonatal male Swiss Webster mice were administered PBS or IL-1β from P1 through P5 and euthanized at P10. The numbers of total cycling cells in the SGZ were assessed using the physical dissector method of stereology. Representative images of Ki67/ DAPI stained 8 μm paraffin sections from P10 (A) control versus (B) IL-1β-treated mice. Scale bar=50 μm. (C) Quantification of the number of Ki67^+^ cells in the SGZ of P10 control versus IL-1β-treated mice. Results are expressed as mean ± SEM; *p<0.05 by unpaired t-test, n=3 mice per group. (D) Timeline of treatment and endpoints for *in vivo* immunofluorescence staining.

To determine if the reduction in cycling cells in the SGZ might be confined to the most rapidly proliferating cells in the SGZ, the TAPs (Figure 3A), mice were treated as previously described and euthanized at P10. Optical fractionator stereology was performed to count the number of cycling TAPs, using Tbr2 and Ki67 as markers for TAPs and cycling cells, respectively, in the SGZ of the dorsal hippocampus. A statistically significant decrease in the number of Tbr2^+^/Ki67^+^ cells (28% decrease) was found in the SGZ of mice receiving IL-1β versus PBS, as determined by unpaired t-test (t(6)=2.513, p=0.046, Figure 3B). To determine if the reduction in cycling TAPs in the SGZ of these mice persisted into adulthood, mice were administered IL-1β or PBS and euthanized at P60. The optical fractionator was used to count the number of cycling TAPs throughout the dorsoventral hippocampus. No statistically significant change in Tbr2^+^/Ki67^+^ cells was found in the SGZ of mice receiving IL-1β versus PBS (unpaired t-test, t(4)=1.06, p=0.35, Figure 3C-D).

**Figure 3.**
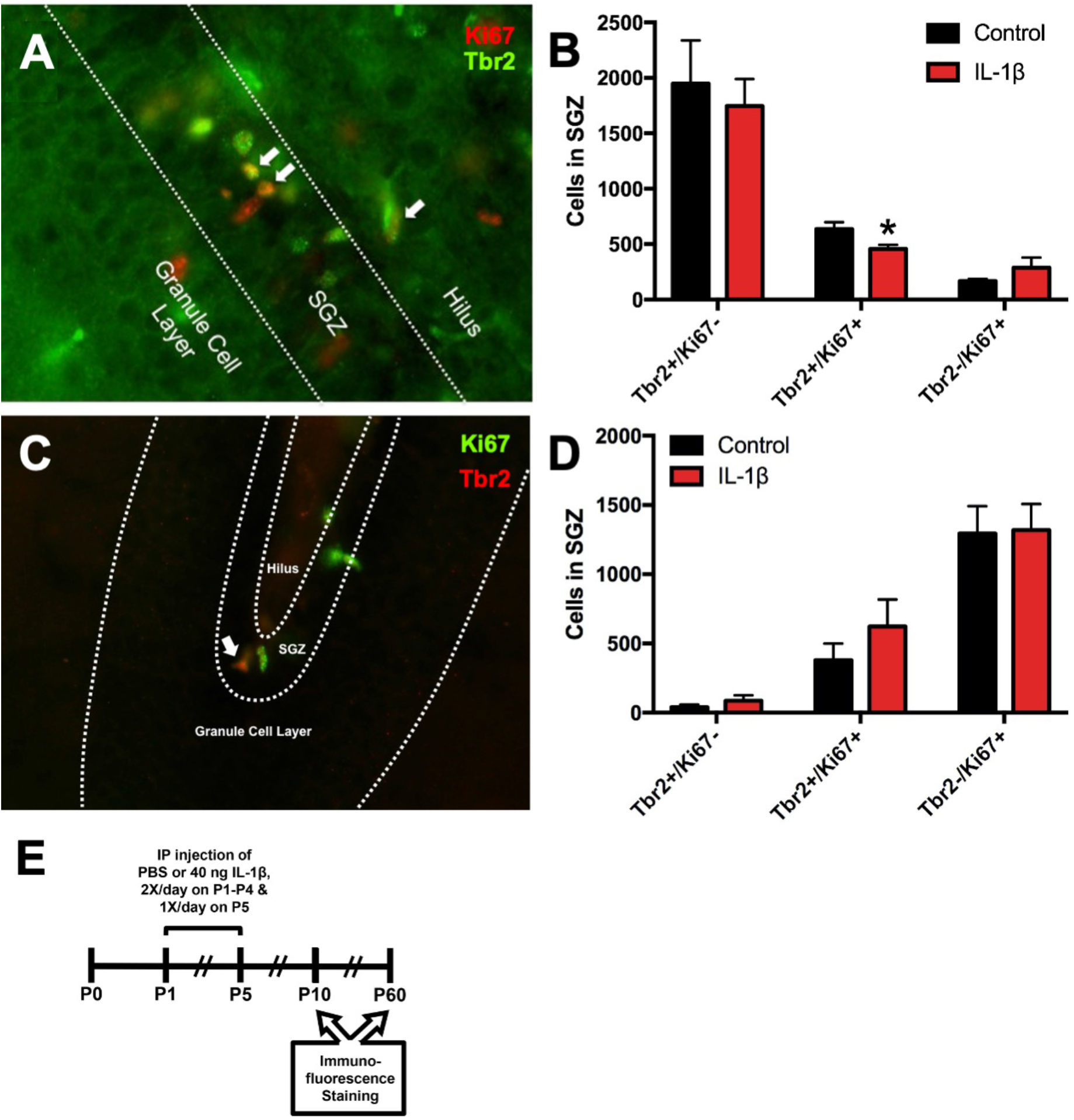
The number of cycling Tbr2^+^ progenitors is acutely reduced in mice with systemic inflammation. Neonatal male Swiss Webster mice were administered PBS or IL-1β from P1 through P5 and euthanized at P10. The numbers of total cycling Tbr2^+^ progenitors in the SGZ were assessed using the optical fractionator method of stereology. (A) Representative of Tbr2/ Ki67 staining of 40 μm sections of fixed-frozen brains from P10 mice. White arrows indicate Tbr2^+^/Ki67^+^ cells. (B) Quantification of the number of Tbr2^+^/Ki67^+^ cells in the SGZ of P10 control versus IL-1β-treated mice. Results are expressed as mean ± SEM; *p<0.05 by unpaired t-test, n=4 mice per group. (C) Representative image of Tbr2/ Ki67 staining of 40 μm sections of fixed-frozen brains from P60 mice. White arrow indicates Tbr2^+^/Ki67^+^ cell. (D) Quantification of the number of Tbr2^+^ and Ki67^+^ cells in the SGZ of P60 control versus IL-1β-treated mice. Results are expressed as mean ± SEM; n=3 mice per group. (E) Timeline of treatment and endpoints for *in vivo* immunofluorescence staining.

### IL-1α and IL-1β reduce the growth of neonatal mouse hippocampal NPs

To evaluate the effect of IL-1 cytokines on neonatal mouse hippocampal NPs, NPs derived from P5 mouse hippocampi were incubated with non-saturating concentrations of IL-1 cytokines (Figure 4A). After propagating primary neurospheres from P5 mouse hippocampi, secondary neurospheres were cultured in the presence of either 50 or 500 pg/mL IL-1β versus control over 6 days (Figure 4E-G). Incubation with 500 pg/mL IL-1β significantly reduced the size of neurospheres by 27% (unpaired t-test, t(4)=3.26, p=0.031) (Figure 4B), but it did not decrease the total number of neurospheres generated (Figure 4C). The reduced neurosphere size was accompanied by a reduced cell yield obtained upon dissociating the spheres (unpaired t-test, t(4)=4.042, p=0.012) (Figure 4D). To determine whether IL-1α has similar effects on neonatal mouse hippocampal NPs, primary neurospheres derived from P5 mouse hippocampi were cultured in the presence of either 500 pg/mL IL-1α or IL-β versus control over 7 days (Figure 4J-L). As expected, the presence of either IL-1β or IL-1α reduced the sizes of neurospheres (Figure 4H). The reduced neurosphere size was also accompanied by a reduced cell yield (Figure 4I).

**Figure 4.**
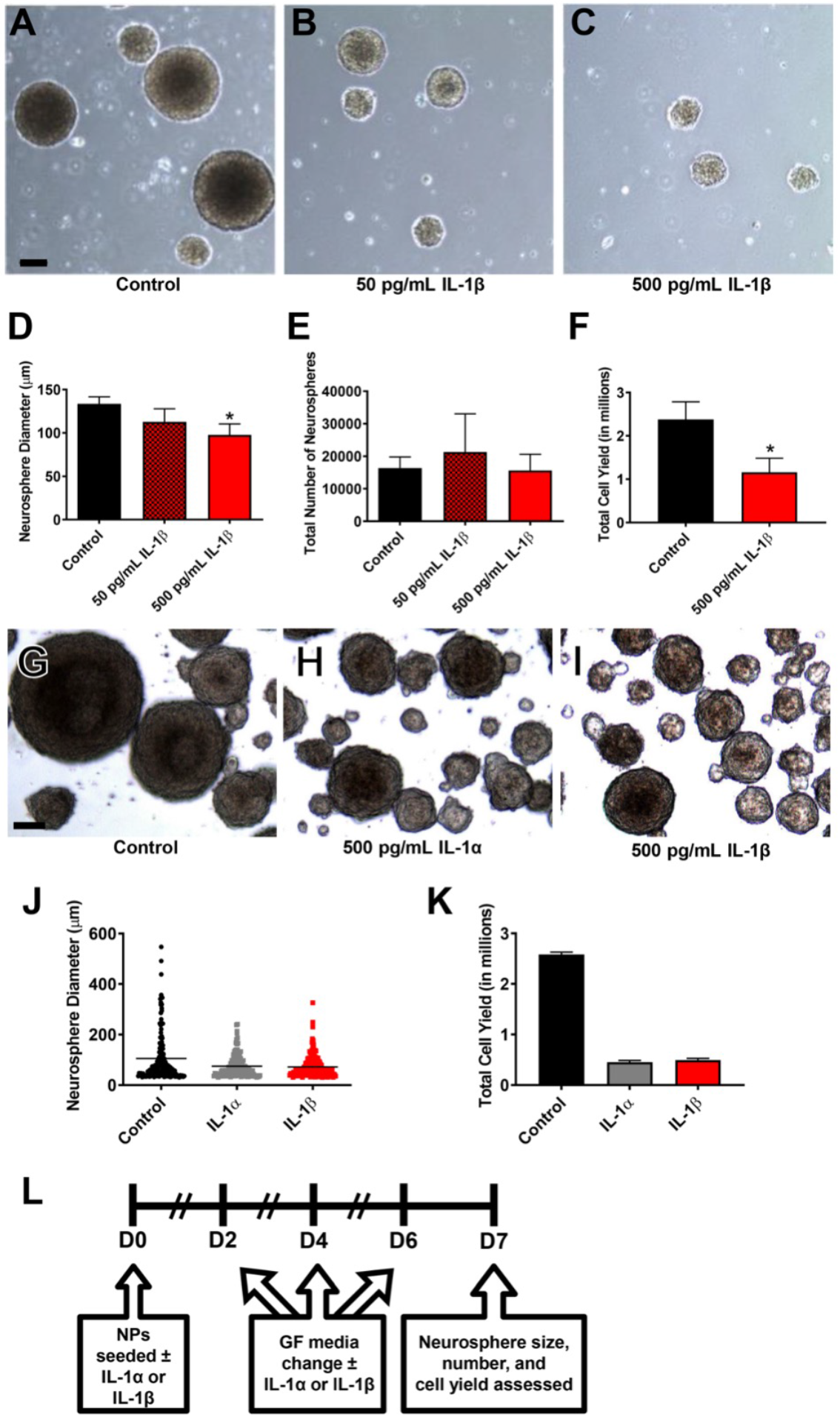
IL-1 treatment reduces the growth of neonatal mouse hippocampal NPs. NPs derived from the hippocampi of neonatal C57BL/6 or Swiss Webster mice were grown in the presence of IL-1 for 7 days, and neurosphere size, counts and cell yield were quantified. Representative images of (A) control versus (B) 50 pg/mL IL-1β and (C) 500 pg/mL IL-1β-treated neurospheres. Quantification of (D) neurosphere size, (E) neurosphere counts, and (F) NP cell yield after 7 days in standard growth medium versus treatment with either 50 or 500 pg/mL IL-1β. Results are expressed as mean ± SEM; *p<0.05 by unpaired t-test. Results are averaged from 3 independent experiments. Scale bar=100 μm. Representative images of (G) control versus (H) 500 pg/mL IL-1α and (I) 500 pg/mL IL-1β-treated neurospheres. Quantification of (J) neurosphere size and (K) total NP cell yield after 7 days in standard growth medium versus treatment with either 500 pg/mL IL-1β or IL-1α. Scale bar=50 μm. Results are expressed as mean ± SEM. Results are averaged from a triplicate experiment. (L) Timeline of treatment and endpoints for neurosphere growth assays.

### IL-1β treatment does not reduce NSC self-renewal or deplete NSCs in neonatal mouse hippocampal NP cultures

To determine whether the negative effect of IL-1β on neonatal mouse hippocampal NP growth was due to an adverse effect on NSC homeostasis, the effect of IL-1β on NSC self-renewal and sphere forming frequency was assessed (Figure 5A). Primary neurospheres were grown in the presence of 500 pg/mL IL-1β versus no treatment and then passaged every 7 days for 3 weeks in the absence of IL-1β, and the cell yield was recorded at each passage. While IL-1β significantly reduced cell yield at passage # 1 when neurospheres were grown in the presence of IL-1β (unpaired t-test, t(4)=3.460, p=0.026), cell yield was not affected at subsequent passages where IL-1β was absent (Figure 5B). To determine whether IL-1β was affecting the sphere forming frequency of the NSCs, primary neurospheres were grown in the presence of 500 pg/mL IL-1β versus no treatment for 7 days. The neurospheres were dissociated and then plated into wells at incrementally fewer cells per well for the limiting dilution analysis (LDA) assay. The LDA assay showed that IL-1β increased the sphere forming frequency (1 in 95 cells) compared to untreated NPs (1 in 189 cells) (Figure 5C). Together, these data suggest that IL-1β does not reduce NSC self-renewal or deplete NSCs. By contrast, IL-1β treatment enriches the proportion of NSCs in the cultures.

**Figure 5.**
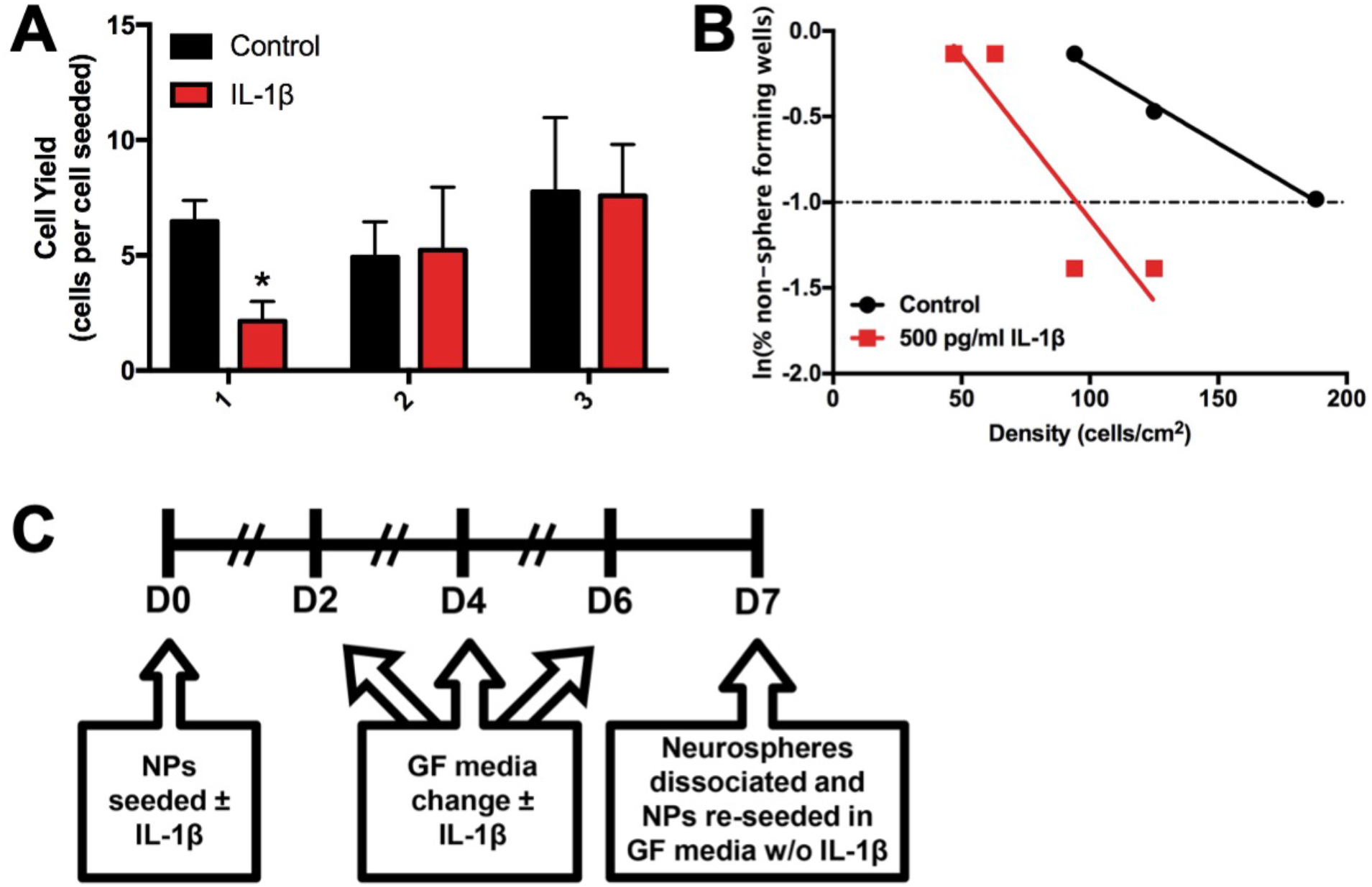
IL-1 treatment does not reduce NSC self-renewal or deplete NSCs in neonatal mouse hippocampal NP cultures. NPs derived from the hippocampi of neonatal Blk6 or Swiss Webster mice were grown in the presence of IL-1 for 7 days and either passaged repeatedly from three weeks or seeded into plates for limiting dilution analysis (LDA). (A) Quantification of cell yield after each passage of NPs that were either maintained in standard growth medium or treated with 500 pg/mL IL-1β. Results are expressed as mean ± SEM; *p<0.05 by unpaired t-test, n=3 independent experiments. (B) Representative graph depicting sphere-forming frequency of control versus 500 pg/mL IL-1β-treated NPs in LDA. (C) Timeline of treatment and protocol for repeated passaging and LDA.

### IL-1α and IL-1β arrest Tbr2^+^ neonatal hippocampal NPs in the G1 phase of the cell cycle

To establish whether the observed effects with IL-1 involved changes in cell cycle progression, neurospheres were starved of EGF and FGF-2 for cell cycle synchronization and allowed to reenter the cell cycle in the presence of either 500 pg/mL IL-1α, IL-1β or PBS. The cells were then processed for a flow cytometric analysis of DNA content in combination with immunofluorescence for the transcription factor Tbr2. As early as 12 hours post-treatment, there was an increase in the percentage of cells in G1 phase and a decrease in the percentage of cells in S phase; however, these values did not achieve statistical significance (data not shown). By 48 hours after treatment, there was a significant 22% and 21% increase in the number of Tbr2^+^ NPs in G1 phase after IL-1α and IL-1β treatment (unpaired t-test, t(4)=2.933, p=0.043) and IL-1β (unpaired t-test, t(4)=3.001, p=0.04) treated spheres versus controls, respectively (Figure 6D). Additionally, significant 41% and 39% reductions in the number of cells in S phase were observed at 48 hours in IL-1α (unpaired t-test, t(4)=8.702, p=0.001) and IL-1β (unpaired t-test, t(4)=8.352, p=0.0011) treated spheres versus controls, respectively (Figure 6D). By Western blot, levels of Cyclin D1 were reduced by 3 h after treatment (Figure 6E), and by 6 h of treatment, levels of Cyclin B/B1 were reduced. These data suggest that IL-1 cytokines reduce the growth of NPs by preventing Tbr2^+^ NP entry into the S phase of the cell cycle by suppressing the requisite increase in Cyclin D1 and Cyclin B in hippocampal NPs.

**Figure 6.**
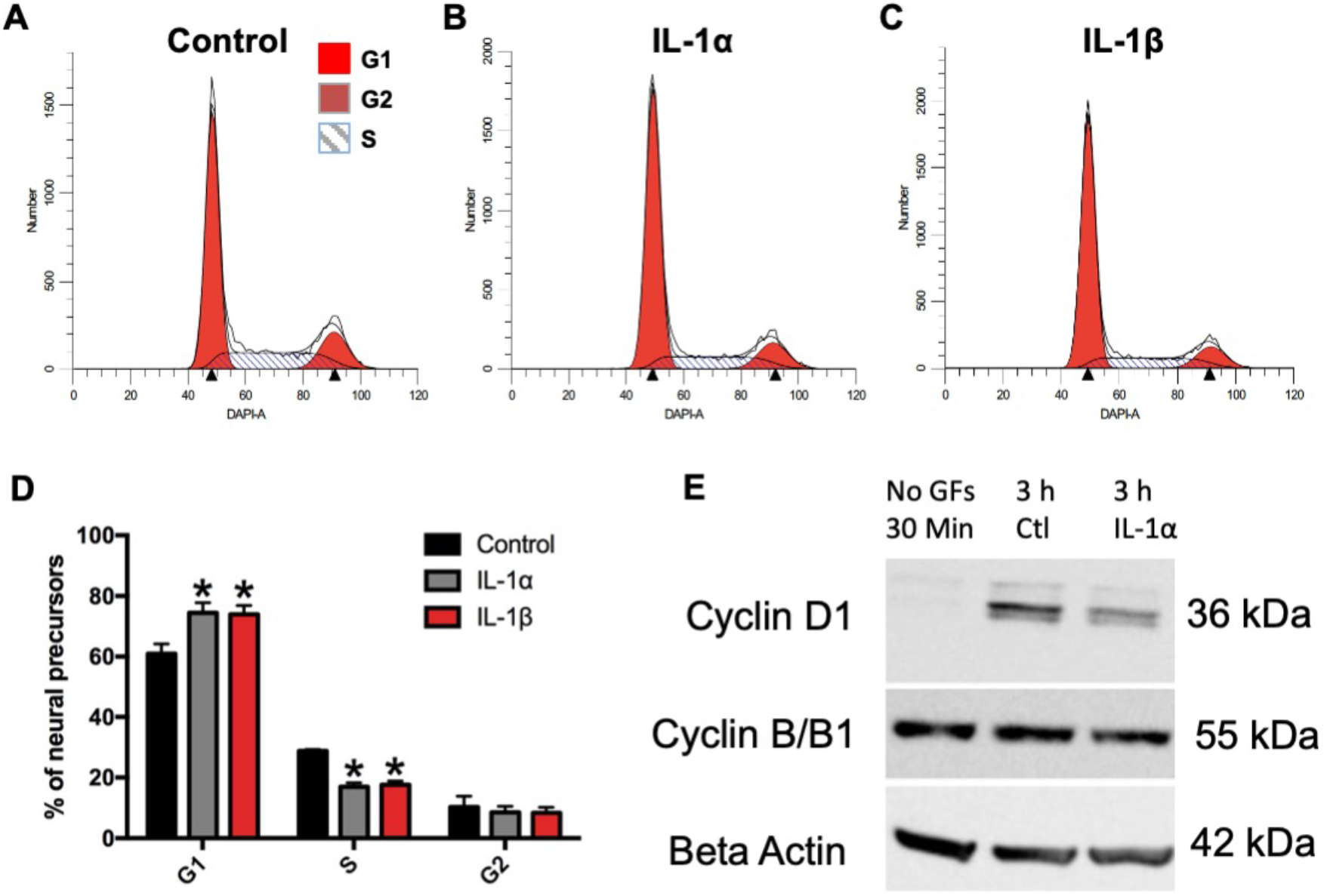
Tbr2^+^ NPs accumulate in G1 due to decreased entry into S phase. NPs were treated with IL-1α or IL-1βfor 48h. Cells were stained for Tbr2 and DAPI and analyzed by flow cytometry. Representative histograms showing cell cycle analysis of (A) untreated NPs versus NPs treated with (B) 500 pg/mL IL-1α and (C) 500 pg/mL IL-1β at 48 hrs post-recovery/ treatment. (D) Percentages of cells in each cell cycle phase were averaged across 3 experiments and expressed as mean ± SEM; *p<0.05 by two-way ANOVA followed by Tukey’s multiple comparisons test. n=3 independent experiments. (E) Analyses of Cyclin D1 and Cyclin B/B1 from untreated NPs versus NPs treated with 500 pg/mL IL-1α at 3 hours post-recovery/ treatment. Data depicted in panel E are representative of 2 independent experiments.

### Mice with systemic inflammation exhibit signs of anxiety and reduced exploratory behavior in adolescence

Neonatal systemic inflammation induced by LPS has been shown to increased anxiety in adult animals(Breivik *et al.*, 2002). To evaluate whether mice with systemic inflammation induced by IL-1β administration exhibited altered anxiety and exploratory behavior, the mice were tested using two different behavioral tasks; elevated plus maze (EPM) at P36 and open field test (OFT) at P39. In agreement with previously published data, IL-1β-treated mice showed increased anxiety-like behaviors indicated by fewer crossings into the open arms (Figure 7A), compared to those treated with PBS (unpaired t-test, t(22)=2.211, p=0.0377). In contrast, there was no difference between total distance travelled (Figure 7B). The open field test at P39 also revealed increased levels of anxiety-like behaviors in IL-1β treated mice, indicated by significantly less time spent exploring the center zone during the first 30 minutes (unpaired t-test, t(22)=2.14, p=0.044) as well as throughout the entire 60 minute test duration (unpaired t-test, t(22)=2.857, p=0.009), compared to PBS treated mice (Figure 7B-D). In contrast, there was no difference between treatments in average speed, total distance travelled, total number of rears and total resting time (Figure 7E,F), indicating that basal locomotor function was not affected.

**Figure 7.**
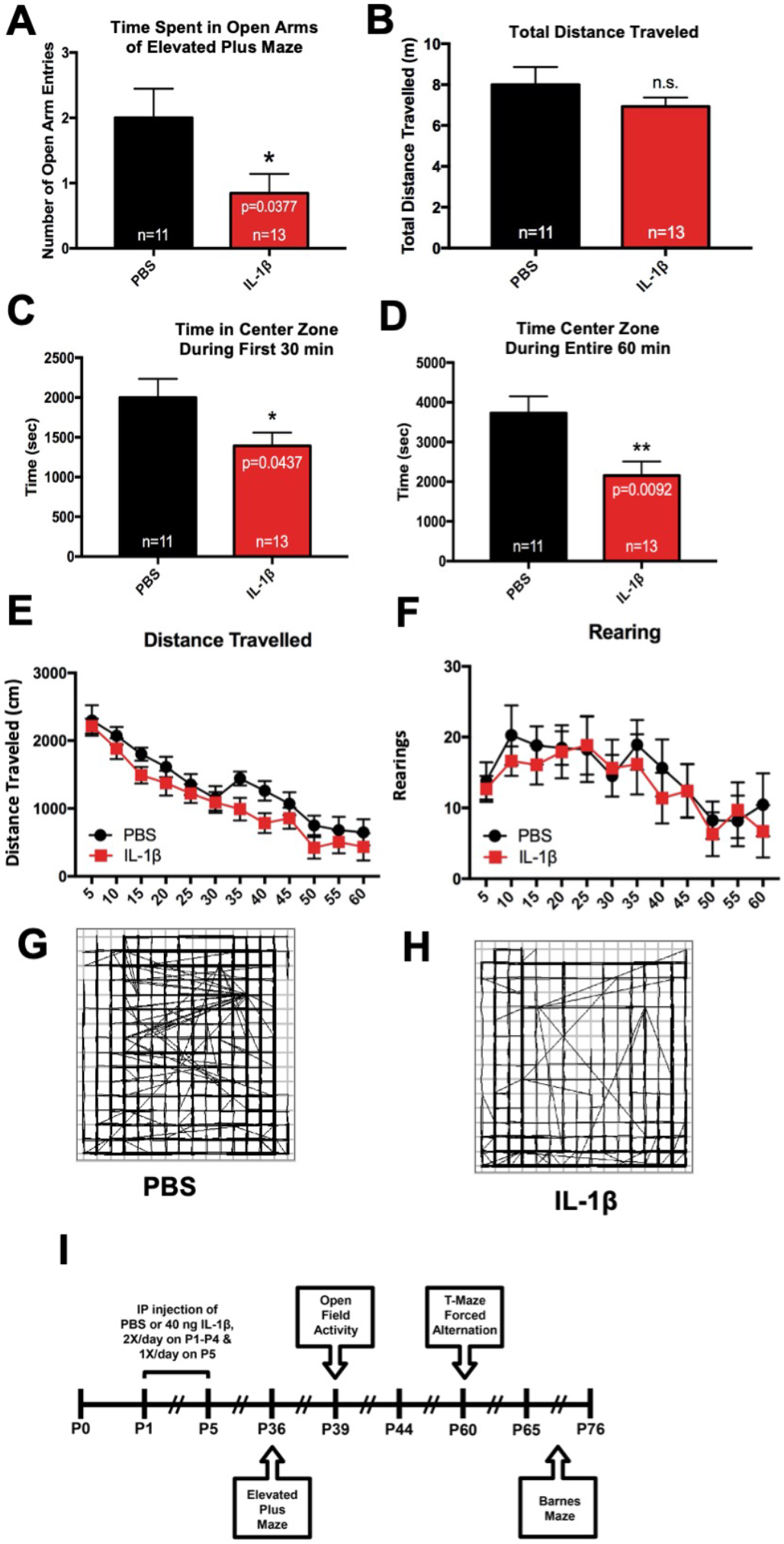
Mice with perinatal systemic inflammation show signs of anxiety in the elevated plus maze and open field activity tasks during adolescence. (A) Neonatal male Swiss Webster mice were administered PBS or IL-1β from P1 through P5 and then tested in the elevated plus maze task for exploratory and anxiety behavior at P36. Results are expressed as mean ± SEM; *p<0.05 by unpaired t-test. (B) Total distance traveled was measured using ANY-maze behavioral tracking software. Mice also were tested using open field activity system for exploratory and anxiety behavior at P39. Quantification of time spent in the center zone of the activity chamber during the (C) first 30 min and (D) the entire 60 min session, (E) total distance traveled and (F) rearing events for control versus IL-1β-treated mice. Results are expressed as mean ± SEM; *p<0.05, **p<0.01 by unpaired t-test, n=11 mice for control group, n=13 mice for IL-1β group. (G and H) Representative path traces for control versus IL-1β-treated mice during a 60 min session. (I) Timeline of treatment and endpoints for these behavioral assessments.

### Mice with systemic inflammation show impairments in long-term spatial reference memory but not shortterm working memory

A previous study showed that male mice that had received systemic administrations of IL-1β from P1 through P5 show reduced performance in working memory tasks at P29 and P30 (Favrais *et al.*, 2011). Furthermore, other studies have found that neonatal inflammation leads to deficits in reference memory in the adulthood (Bilbo *et al.*, 2005; Harre *et al.*, 2008). To determine whether perinatal IL-1β treatment leads to deficits in working and reference memory in early adulthood, two different behavioral tasks were used. First, we used the T-maze forced alternation task at P60 to test for spatial working memory in IL-1β and PBS treated mice. Interestingly, mice treated with IL-1β displayed significantly higher rates of alternation compared to those treated with PBS (unpaired t-test, t(20)=2.481, p=0.022), indicating that working memory was enhanced in IL-1β-treated mice (Figure 8A).

**Figure 8.**
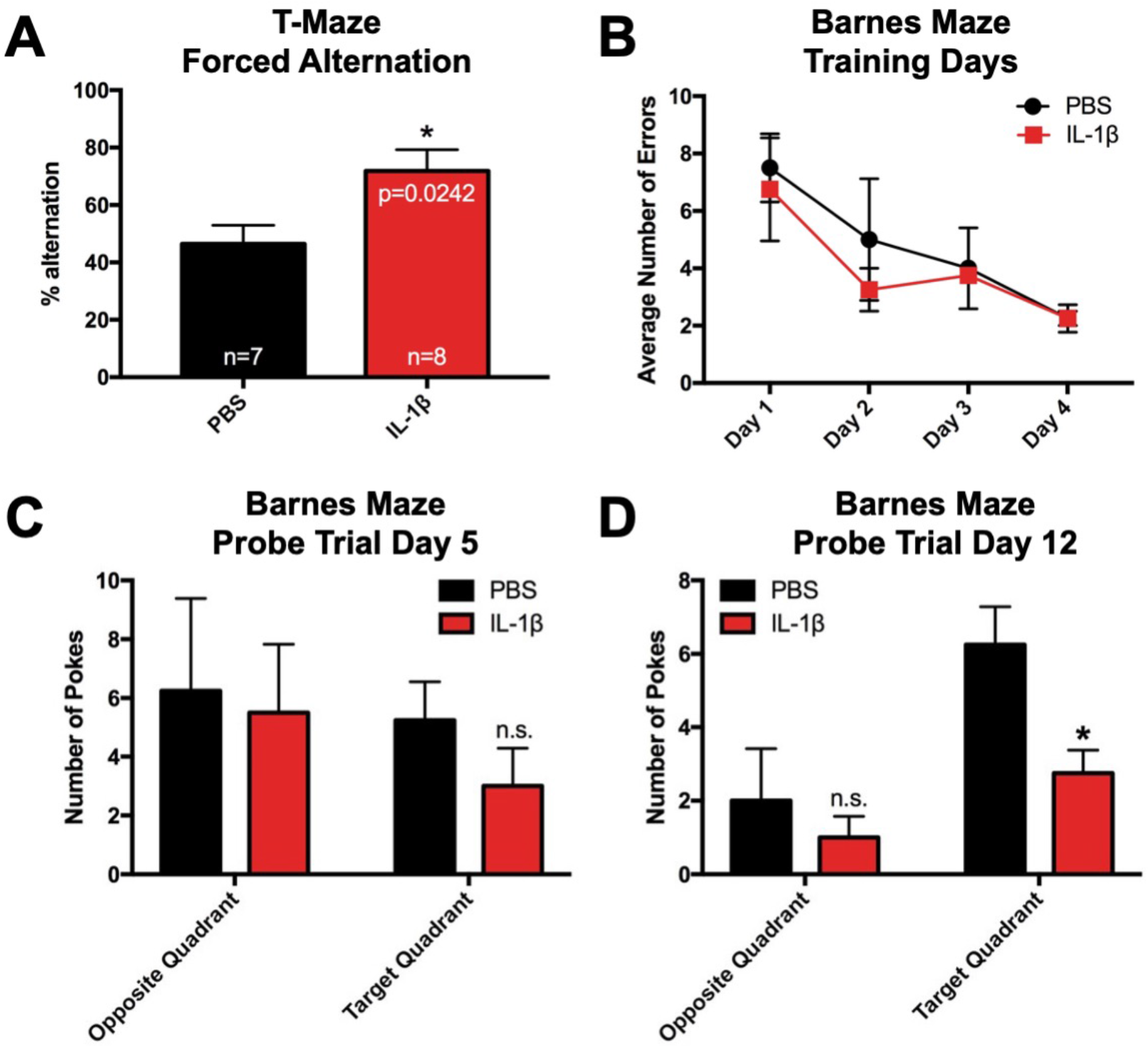
Mice with perinatal systemic inflammation show enhanced short-term working memory and deficits in long-term spatial memory in early adulthood. (A) Neonatal male Swiss Webster mice were administered PBS or IL-1β from P1 through P5 and then tested in the T-maze forced alternation task at P60. Quantification of the percent alternation of control versus IL-1β-treated mice in the T-maze forced alternation task. Results are expressed as mean ± SEM; *p<0.05 by unpaired t-test, n=7 mice for control group, n=8 mice for IL-1β group. Mice also were tested in the Barnes maze task at P65 through P76. (B) Average number of errors on training Days 1 through 4 of control versus IL-1β treated mice. Average number of head pokes into the target quadrant versus the opposite quadrant of the Barnes maze during the 90 sec probe trial on (C) Day 5 and (D) Day 12. Results are expressed as mean ± SEM; *p<0.05 by unpaired t-test

To assess the impact of perinatal IL-1β on spatial memory function after TBI, IL-1β- and PBS-treated mice were tested using the Barnes maze task with visual cues, where they learned the location of the escape hole over 4 consecutive days (P65~P68), followed two probe trials on P69 and P76, where the escape box was removed and the mice were allowed to search for its prior location (Vorhees Cv Fau – Williams and Williams). IL-1β- and PBS-treated mice learned the task at the same rate (Figure 8B). Subsequently, the first probe trial at P69 revealed no sign of reference memory deficits (Figure 8C). In contrast, the second probe trial at the longer time point of P76 revealed a significant reduction in the average number of head pokes in target quadrant (unpaired t-test, t(6)=2.898, p=0.027), which is indicative of a reference memory deficit (Figure 8D). Altogether, these data demonstrate that that systemic inflammation induced by perinatal IL-1β leads to a persistent long-term reference memory deficit.

## Discussion

The results of these studies demonstrate that IL-1β-mediated systemic inflammation in neonatal mice during the first 5 days of life increases tissue levels of the pro-inflammatory cytokines and chemokines IL-1α, CXCL1,CCL2 and VEGF-A acutely within the hippocampus. The increases in the levels of these cytokines was accompanied by a decrease in the proliferation of the Tbr2^+^ intermediate progenitors of the DG. Taking into consideration the high level of expression of the IL-1 receptor within the immature DG of the hippocampus we further explored the role of IL-1 signaling on the stem cells and progenitors of the DG (Cunningham *et al.*, 1992). Our *in vitro* analyses demonstrate that either IL-1α or IL-1β at non-receptor-saturating concentrations reduce the number of cycling Tbr2^+^ TAPs by suppressing the expression of requisite cell cycle proteins Cyclin D1 and Cyclin B, thus preventing the progenitors from entering S phase. As a consequence, IL-1 enriched NP cultures for NSCs during cytokine exposure; however, NPs regained their proliferative capacity upon terminating IL-1 exposure. While *in vivo* neurogenesis eventually recovered from exposure to perinatal systemic inflammation, changes in hippocampal dependent behaviors were persistently affected, including increased signs of anxiety and long-term spatial memory deficits. Altogether, these data support the view that perinatal inflammation negatively affects the developing hippocampus, resulting in behavioral deficits that persist into adulthood. These data provide a new perspective into the origin of the cognitive and behavioral impairments observed in prematurely born sick infants.

Our studies extend the body of evidence that indicate that perinatal inflammation adversely affects the developing brain. Notably, our data show the hippocampus, which is a late developing structure is affected by systemic inflammation occurring at the human equivalent of the beginning of the 3rd trimester. Within the adult mouse hippocampus, approximately 75% of proliferating cells in the DG are Tbr2^+^ (Hodge *et al.*, 2008). These Tbr2^+^ cells are progenitors for the glutamatergic neurons of the DG. Therefore, any effects on the dynamics of Tbr2^+^ cell proliferation could lead to abnormal development of the DG, which continues to grow for three weeks postnatally in the mouse (Chuang *et al.*, 2011). Previous *in vivo* studies have reported acute negative effects on the NPs of the SGZ in response to perinatal inflammation. However, conflicting results have been reported for long-term effects. In a rat model for perinatal systemic inflammation, LPS administered during late gestation (G20 through G22-23) resulted in a decrease in Dcx labeled cells at P40, suggesting a persistent effect on the NPs of the SGZ (Girard *et al.*, 2012). However, in another study by Smith et al. 2014, where a single dose of LPS was administered at P5, there was only a transient decrease in Dcx labeled cells in the SGZ at P21 and no long-term-effect on neuronal survival when analyzed at P74 (Smith *et al.*, 2014). Our data using IL-1β to drive neonatal systemic inflammation mirrors the effect seen by Smith et al. in that the effects of systemic inflammation on the NP pool is transient. The differences between our studies and studies reporting sustained reduction in neurogenesis in adulthood are likely due to the developmental window during which the inflammatory response was initiated in each respective model. In the adult mouse, the conditional knock out of MyD88 in Nestin^+^ cells does not abrogate the negative effects on neurogenesis due to the constitutive expression of IL-1β achieved with an adeno-associated viral vector serotype 2 expressing a IL-1β under a CMV promoter (Wu *et al.*, 2013). However, in a rat model of perinatal systemic inflammation induced with IP injections of LPS, negative effects on neurogenesis were abrogated with IL-1Ra (Girard *et al.*, 2012). These results suggest that the Nestin^+^ NPs (i.e., the NSCs) are not adversely affected by increased levels of IL-1, but that the reduced neurogenesis is due to an effect upon the intermediate progenitors (i.e., Tbr2^+^ NPs). Similar results have recently been reported in a study where 1 mg/kg LPS was administered to young adult rats (Melo-Salas *et al.*, 2018).

Many studies on brain insults and injury to the hippocampus have focused on the effects of IL-6 and IL-1β; however, our data show increased expression of IL-1α within the hippocampus. Levels of IL-6 and IL-1β were unchanged within the hippocampus, both at the mRNA and protein level subsequent to IL-1 treatment, suggesting that the negative effects of neuroinflammation in this model is driven by the production of IL-1α. These IL-6 and IL-1β findings are in agreement with previous microarray studies from this model in the forebrain at P5 and P10, where both IL-1β and IL6 were not significantly increased in total homogenates, although IL1a was also not increased. However, the dentate gyrus may be more vulnerable to injury as suggested by localized (and subtle) injury to the blood brain barrier only in the hippocampus in this model (Krishnan *et al.*, 2017)).

Cells of monocytic lineage, specifically macrophages and dendritic cells, are major producers of IL-1α peripherally (Dinarello, 1996). Additionally, studies have shown that platelets are major producers of IL-1α in ischemic brain injury and that platelet derived IL-1α drives cerebral inflammation by activating the cerebrovascular endothelium (Thornton *et al.*, 2010); this is of particular importance since the hippocampus is a highly vascularized brain structure and relatively susceptible to changes in the blood-brain barrier. In models of ischemic brain injury (Denes *et al.*, 2011) microglial production of IL-1α precedes the production of IL-1β. The brain endothelium is also activated by IL-β via the MAPK and NF-κB signaling pathways, resulting in the expression of cell adhesion molecules and various chemokines that aid in leukocyte infiltration (Thornton *et al.*, 2010).

While activated microglia are major producers of IL-1 cytokines in the inflamed brain, studies show that astrocytes are also important producers of IL-1 in brain inflammation. In a mouse model of perinatal systemic inflammation induced by LPS, parenchymal and vascular astrogliosis and parenchymal microgliosis are seen acutely and followed by vascular microgliosis, suggesting that inflammation at the blood brain barrier is first mediated by astrocytic activation followed by microglial activation (Cardoso *et al.*, 2015). Another study supports an early role for astrocytes in chronic brain inflammation. Garber et al., 2018 reported reduced neurogenesis accompanied by astrogliogenesis at 6 days post-infection with West Nile virus, and at 25 days-post infection, astrocytes were found to be the major producers of IL-1β mRNA transcripts compared to microglia. Additionally, they showed that neurogenesis recovered more rapidly in IL-1R1 knock-out mice and was accompanied by reduced astrogliogenesis. Furthermore, they showed that cognition was also recovered in the IL-1R1 knock-out mice, as measured by performance in the Barnes maze task at days 46 through 50 post-infection (Garber *et al.*, 2018). While they attributed these effects to IL-1β, their studies did not rule out a role for IL-1α.

While we observed increases in IL-1α cytokine levels at P10 in mice with systemic inflammation, we also observed increases in CXCL1, CCL2, and VEGF-A proteins. CXCL1 is recognized for its role in neutrophil chemotaxis and CCL2 is known for its roles in monocyte and basophil chemotaxis. The upregulation of these ligands suggests that there is an active neuroinflammatory response that is recruiting neutrophils, monocytes, and basophils to the hippocampus. A recent study that used this model to elucidate the mechanisms responsible for the dysgenesis of the white matter in this model of Encephalopathy of Prematurity demonstrated that microglia become activated within hours of IL-1β administration, and that this activation is due to decreased WNT signaling (reduced ligands and receptors) specifically in the microglia (Van Steenwinckel *et al.*, 2019). We did not evaluate levels of WNTs in the hippocampus as previous studies have shown that WNT signaling increases hippocampal DG neurogenesis. For example, a study by Choe and Pleasure 2012 showed that forced WNT signaling in DG NSCs increased the numbers of intermediate progenitors, which is opposite to the observed effects of IL-1 ligands (Choe and Pleasure).

VEGF-A is recognized for its role as an inducer of angiogenesis and is also known for promoting neurogenesis. Studies from Jin et al., 2002 showed that VEGF-A stimulates the proliferation of immature neurons *in vitro* and that intracerebroventricularly administered VEGF-A increased proliferation of neuronal, astroglial, and endothelial cells 7 days after treatment (Jin *et al.*, 2002). Similarly, increased NP proliferation seen after TBI has been attributed to increased levels of VEGF (Neuberger *et al.*, 2017). These studies suggest that the VEGF is not responsible for the decreased cell proliferation that we observed. However, it is possible that the increased VEGF-A seen at P5 and P10 may be contributing to the restoration of proliferating Tbr2^+^ progenitors observed at P60 in our studies.

In previous studies performed in our lab by Covey et al., 2011, the COX-2 inhibitor, NS398 abrogated the increased proliferation of neonatal-derived SVZ NPs observed in the presence of 5 ng/mL IL-6, suggesting that the IL-6 mediated effects on NP growth were indirect and likely mediated by prostaglandins produced via COX-2 (Covey *et al.*, 2011). A role for COX2 in driving injury to the brain in this model is also supported by our findings that treatment with the COX inhibitor nimesulide prevented the systemic inflammation-induced white matter injury (Shiow *et al.*, 2017). To test whether prostaglandins produced by COX-2 might be mediating the effects of IL-1 on hippocampal NPs, we evaluated whether the addition of NS398 could abrogate the negative effects of IL-1 on NPs expansion. Contrary to our expectations hippocampal NP proliferation was not reduced with NS398, supporting the conclusion that IL-1 directly affects hippocampal NPs (Figure S2).

Our data show that at non-saturating concentrations, IL-1 negatively regulates the proliferation of Tbr2^+^ NPs via G1 to S phase cell cycle arrest, but it does not result in cell death (see Figure S3) or depletion of NSCs *in vitro,* which support previous studies performed in rat cultures (Crampton *et al.*, 2012; Green *et al.*, 2012; Koo and Duman, 2008). The observed cell cycle arrest likely accounts for the reduction in the size of the neurospheres with IL-1 cytokines. Our *in vitro* results further suggest that the IL-1 cytokines preferentially affect the highly proliferative Tbr2^+^ cells without adversely affecting the NSCs. These results are in contrast to studies of Koo and Duman, 2008, which concluded that IL-1 reduced the proliferation of the NSCs (Koo and Duman, 2008), using only Nestin as a marker for the NSCs. It is now generally accepted that cell cultures are comprised of a heterogenous population of cells containing both NSCs and less primitive NPs. If it were the case that the proliferation of NSCs was affected, fewer neurospheres would have formed in our studies where NPs were grown in the presence of IL-1 cytokines. However, the number of neurospheres was not different between control and IL-1 treated group. In fact, as revealed by the LDA experiment, NSCs were enriched in IL-1 treated cultures, suggesting that NSCs maintained their self-renewal capacity after exposure to IL-1 cytokines. While these findings are in contrast with the findings of Koo and Duman, they are supported by studies by Wu et al., 2013 that showed that reduced neurogenesis in the SGZ was not due to IL-1 signaling in the Nestin^+^ population of NPs (Wu *et al.*, 2013). Similarly, a study by Kokovay et al. 2012 showed that treating neurospheres derived from the adult mouse SVZ with 10 ng/mL IL-1β overnight significantly increased the expression of VCAM1 on GFAP^+^ NPs as well as the number of GFAP^+^VCAM1^+^ NPs (Kokovay *et al.*, 2012). These results were accompanied by a significant increase in the number of CD133^+^ cells and a decrease in the number of Ki67^+^ and Mash1^+^ cells, suggesting that IL-1β favored promoted the expansion of NSCs. Other reports on the effects of IL-1β on fetal mouse NPs *in vitro* suggest that IL-1β causes cell cycle arrest and apoptosis via a p53 dependent mechanism; however these studies were performed with 50 ng/mL IL-1β, a supraphysiological concentration of IL-1β that is not likely to mirror the *in vivo* situation (Guadagno *et al.*, 2015).

To determine which cell cycle proteins might be involved in the G1/S cell cycle arrest observed in hippocampal NPs with IL-1 treatment, the levels of cyclin-dependent kinase inhibitors from the CIP/KIP family, which include p21, p27, and p57, were assessed. However, no consistent differences in the levels of KIP proteins or p53 were seen at any timepoint during an 18-hour cell cycle study (Figure S4). By contrast, levels of 2 key cyclins (Cyclin D1 and Cyclin B/B1) were suppressed by IL-1β treatment. Our Western blot results show reduced levels of Cyclin D1 in NPs *in vitro* early in the cell cycle with IL-1α treatment. Interestingly, our cell cycle results also show an accumulation of cells in G0/G1 accompanied by a reduction in the number of cells in G2/M with IL-1 treatment. Our Western blot results also showed reduced levels of Cyclin B/B1 later in the cell cycle with IL-1 treatment. Taken together, the Cyclin D1 and Cyclin B/B1 Western blots suggest that IL-1 exerts cell cycle control of NPs at the G1/S phase transition. These data correlate well with *in vitro* studies that have shown that IL-1β signaling negatively affects rat embryonic hippocampal neurogenesis by decreasing the proliferation of NPs and suppressing neuronal differentiation (Green *et al.*, 2012), via activation of NF-κB signaling (Koo and Duman, 2008; O’Leime *et al.*, 2018). Interestingly, IL-1β suppressed the expression of the orphan nuclear receptor tailless homolog (TLX), that promotes NP proliferation and inhibits neuronal differentiation. Lentiviral overexpression of TLX counteracted the negative effects of IL-1β on the proliferation of embryonic rat NPs *in vitro* (O’Leime *et al.*, 2018).

Dentate granule neurogenesis contributes to the acquisition of contextual memory (Jessberger *et al.*, 2009). Therefore, it is plausible that short- or long-term changes in neurogenic output of the dentate granule layer may affect contextual memory. Indeed, IL-1β-treated neonatal mice display deficits in object location memory and novel object recognition tasks, suggesting that contextual memory is negatively affected by systemic inflammation (Favrais *et al.*, 2011). The results of the additional behavioral tests that we have employed further show that neonatal IL-1 exposure results in persistent deficits in cognition and affect. The novel object recognition task involves the perirhinal cortex and the hippocampus in cases when recency memory or temporal order memory is needed for the task (Barker *et al.*, 2007; Barker and Warburton, 2011). By contrast, the object location memory task involves the perirhinal cortex, the medial prefrontal cortex and the hippocampus, which plays a role in spatial context for the task (Barker and Warburton, 2015).

Considering the regions on the brain involved with these tasks, it was conceivable that poor performance in both of these tasks could be due abnormal function of the perirhinal cortex and/or the hippocampus. Taking into account that the brain is rapidly growing in the neonatal rodent, and correlatively in the third trimester human fetus, changes in the development of any of these structures could lead to abnormal behavior in these tasks. Our results support the conclusion that altered hippocampal developmental is responsible for the deficits in spatial memory and that this hippocampal abnormality will affect adult hippocampal function in long-term memory. By contrast our results demonstrate that there are no deficits in working memory and even suggest enhanced working memory function, as detected in the T-maze forced alternation task. As the goal of this study was to evaluate inflammation induced changes to the hippocampus, we did not evaluate other brain structures; however, a testable hypothesis is that the improvement in working memory is due to alterations in either the production of interneurons within the cingulate cortex or amygdala or in the balance of excitatory/inhibitory synapses in these structures(Bernier *et al.*, 2002; Inta *et al.*, 2008) resulting in an increased state of arousal that improved performance on the forced alternation task (Bernier *et al.*, 2002; Inta *et al.*, 2008). The enhanced memory might be attributed to the effects of low concentrations of IL-1β on normal memory function (Desson and Ferguson, 2003; Murray *et al*.).

Additionally, our results demonstrate neonatal IL-1β exposure increases anxiety that appears during adolescence and lasts throughout adulthood. Since affective behaviors involve multiple regions of the brain that are connected to the DG both directly (i.e., the locus coeruleus), and indirectly (i.e., the amygdala), is it conceivable that changes in either of these regions or the hippocampus during development could lead to an anxiety phenotype, which are subjects for future studies.

To summarize, these studies show that systemic inflammation induced by repeated administration of IL-1β during the first 5 days of life increased hippocampal levels of IL-1α and acutely reduced the proliferation of Tbr2^+^ NPs in the SGZ of mice. *In vitro,* both IL-1α and IL-1β produced G1/S cell cycle arrest that resulted in reduced NP proliferation within the TAP cell cohort due to decreased expression of essential cyclins. By contrast, IL-1β treatment increased NSC frequency. Upon terminating IL-1β treatment, the progenitor cell pool regained its proliferative capacity. *In vivo* studies show that mice that received IL-1β as neonates were more anxious in several behavioral assays during adolescence that persisted into adulthood. Furthermore, these mice did not display short-term working memory deficits in adulthood, but they did display deficits in long-term spatial memory. Altogether, our data support the view that perinatal inflammation negatively affects the developing hippocampus, producing behavioral problems that persist into adulthood. These data provide a new perspective into the origin of the cognitive and behavioral impairments observed in prematurely-born sick infants.

## Acknowledgements

The authors would like to thank Leslie Schwendimann and Dr. Sophie Lebon for the advice and assistance they provided to Dr. Levison while he was on sabbatical in Dr. Gressens’ laboratory. A preliminary report of this work was presented at the 9^th^ Hershey Conference on Developmental Brain Injury at St. Michaels, MD, June 6, 2014 and at the Fusion Neurogenesis Conference in Cancun, MX, March 2016.

## Supplemental Figures

**Figure S1.**
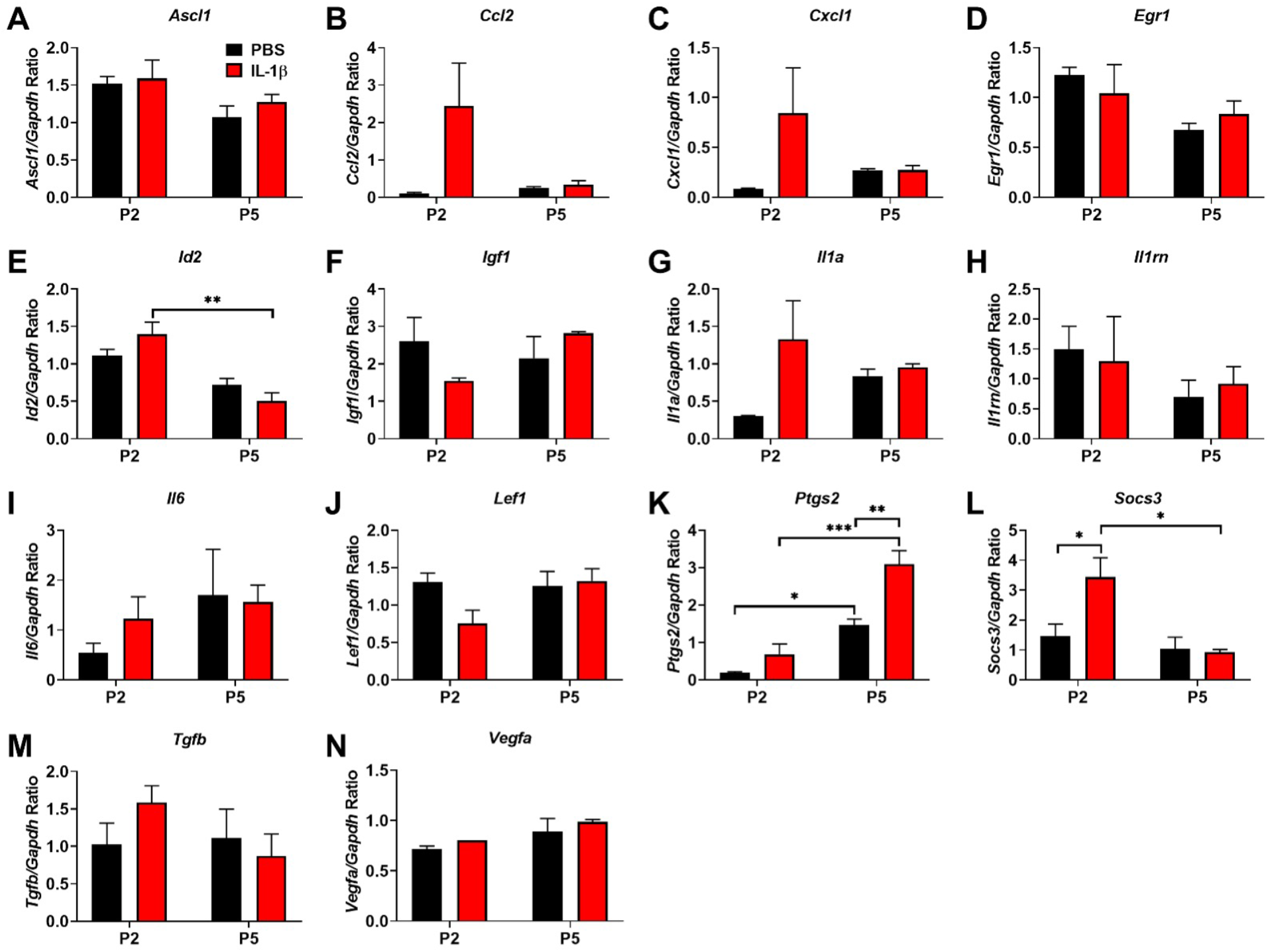
Ptgs2 and Socs3 mRNA levels are significantly elevated acutely in the hippocampus of mice with systemic inflammation. Neonatal male Swiss Webster mice were administered PBS or IL-1β from P1 through P5 and euthanized at P2 and P5. Hippocampal mRNA was extracted and expression levels of transcripts for (A) *Ascl1,* (B) *Ccl2,* (C) *Cxcl1,* (D) *Egr1,* (E) *Id2,* (F) *Igf1,* (G) *Il1a,* (H) *Il1rn,* (I) *Il6,* (J) *Lef1,* (K) *Ptgs2,* (L) *Socs3,* (M) *Tgfb,* and (N) *Vegfa* were assessed using qRT-PCR normalized to GAPDH. Results are expressed as mean ± SEM; *p<0.05, **p<0.01, and ***p<0.001 by two-way ANOVA with Tukey’s multiple comparisons test, n=5 mice per group.

**Figure S2.**
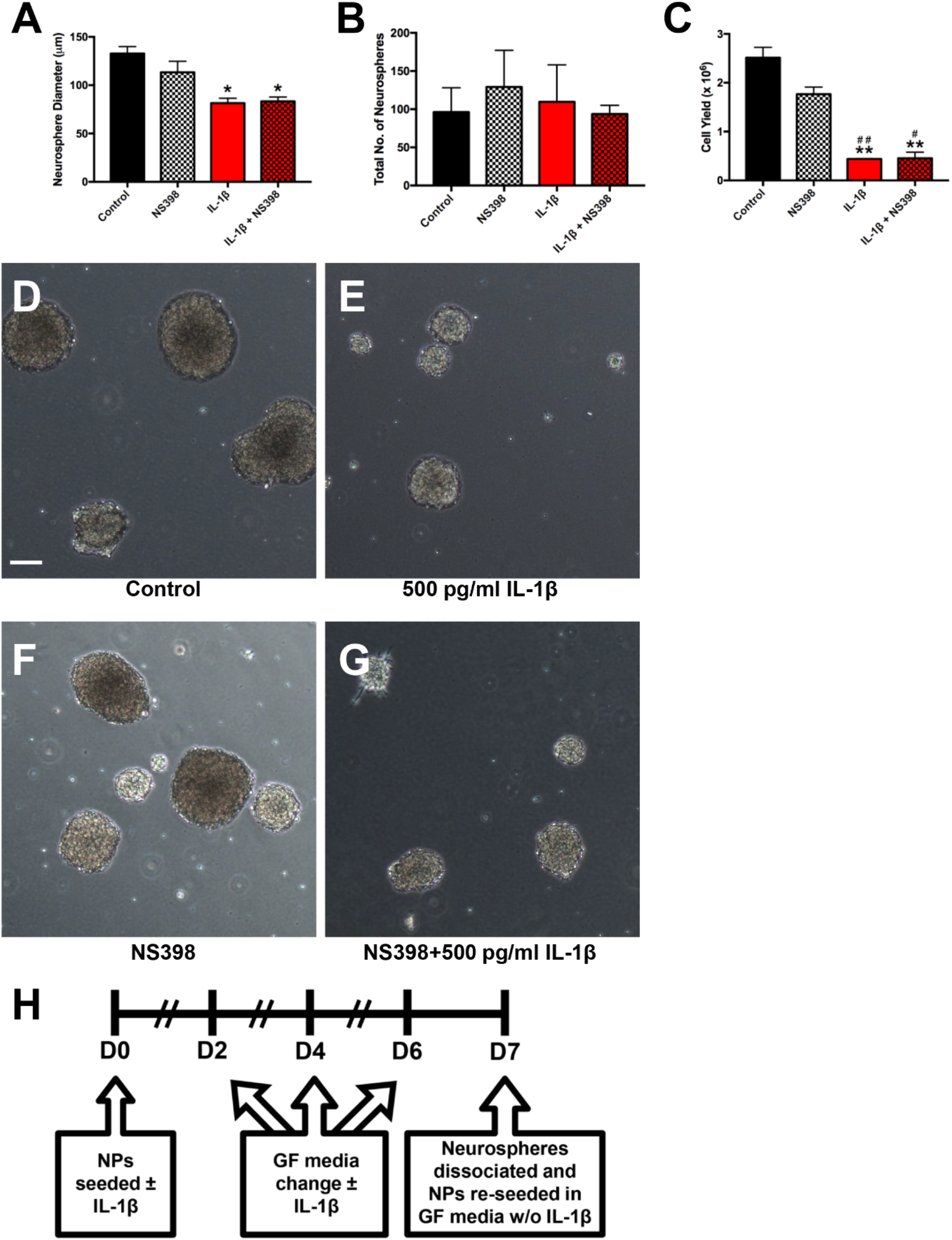
The Cox-2 inhibitor, NS398, does not rescue IL-1-mediated reduction in growth of neonatal mouse hippocampal NPs. The *in vivo* study evaluating the effect of systemic administration of IL-1β on the hippocampal mRNA expression of cytokines and other proteins involved in inflammation revealed that *Ptgs* was significantly increased in both control and IL-1β-treated pups at P5 compared to P2 but that significantly higher expression of *Ptgs* was observed in IL-β-treated pups compared to control at P5 (Figure S1); these data suggested that IL-1 might be exerting its effects through the production of prostaglandins. Therefore, the ability of the small molecule Cox-2 inhibitor, NS398, to rescue neonatal mouse hippocampal NPs from the effects of IL-1 was investigated. NPs derived from P5 mouse hippocampus were incubated with 500 pg/mL IL-1β in the absence or presence of 2 μg/mL NS398 (Figure S2H). A one-way ANOVA revealed significant differences among group means for neurosphere diameter (F(3,4)=10.72, p=0.0221; Figure S2A). Tukey’s multiple comparisons test detected a significant 40% decrease in neurosphere diameter with treatment of IL-1β alone compared to untreated (q(4)= 6.779, p<0.05). Similarly, a significant 37% decrease in neurosphere diameter with treatment of IL-1β plus NS398 compared to untreated was detected (q(4)= 6.523, p<0.05). Additionally, a one-way ANOVA revealed significant differences among group means for cell yield (F(3,4)=50.89, p=0.0012; Figure S2C). Tukey’s multiple comparisons test detected a significant 83% decrease in cell yield with treatment of IL-1β alone compared to untreated (q(4)= 14.46, p<0.01) and a significant 75% decrease in cell yield with treatment of IL-1β alone compared to NS398 alone (q(4)=9.25, p<0.01). Similarly, a significant 82% decrease in cell yield with treatment of IL-1β plus NS398 compared to untreated was detected (q(4)= 14.34, p<0.01) and a significant 74% decrease in cell yield with treatment of IL-1β plus NS398 compared to NS398 alone (q(4)=9.135, p<0.05). No significant differences were found between the untreated and NS398 alone treatment with regard to neurosphere diameter or cell yield. Additionally, no changes among group means for neurosphere number were detected with one-way ANOVA (Figure S2B). Together, these data show that Cox-2 inhibition with NS398 alone does not alter normal neurosphere growth and that Cox-2 inhibition with NS398 does not rescue the reduced neurosphere growth attributed to IL-1 treatment.

**Figure S3.**
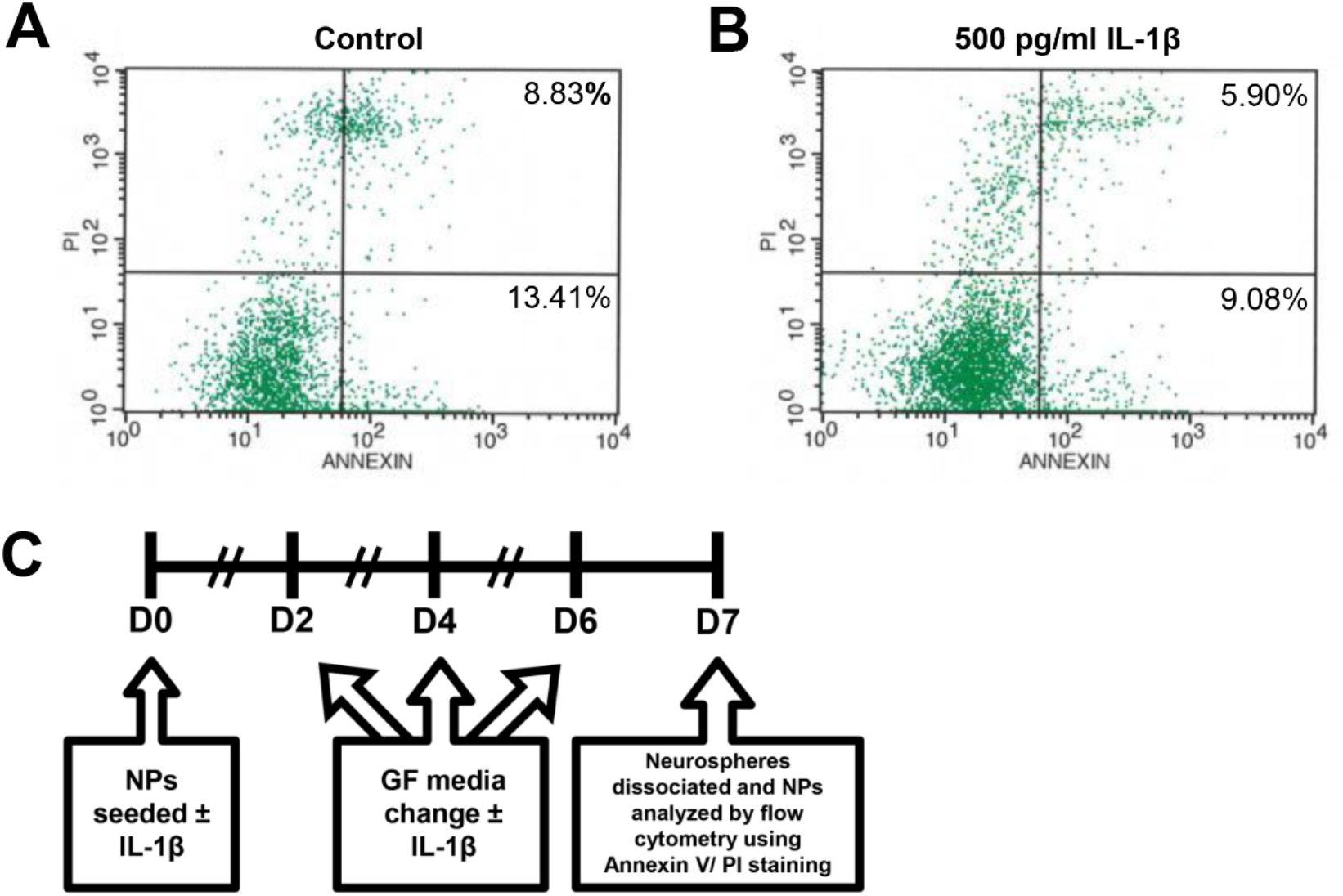
IL-1 treatment does not induce cell death in neonatal mouse hippocampal NPs. The decrease in the number of cells per sphere resulting from NP incubation with IL-1 could arise because of reduced cell proliferation or increased cell death. To evaluate whether the reduction in growth of neonatal mouse hippocampal NPs was due to cell death, secondary neurospheres were cultured in the presence of 500 pg/mL IL-1β versus control over 7 days, after which neurospheres were dissociated and assessed for cell death by staining for cell surface Annexin V and the vital dye propidium iodide, PI, and analyzed by flow cytometry (Figure S3C). Cells categorized as in the early stages of cell death were Annexin V^+^, whereas cells in the late stage of apoptosis were characterized as Annexin V^+^/PI^+^. Negating the conclusion that IL-1 treatment was inducing apoptotic cell death, there was no increase in either the percentage of early or late apoptotic cells with IL-1β treatment versus control (Figure S3A-B). These data suggest that the reduction in NP growth resulting from incubation with IL-1 is not due to cell death.

**Figure S4.**
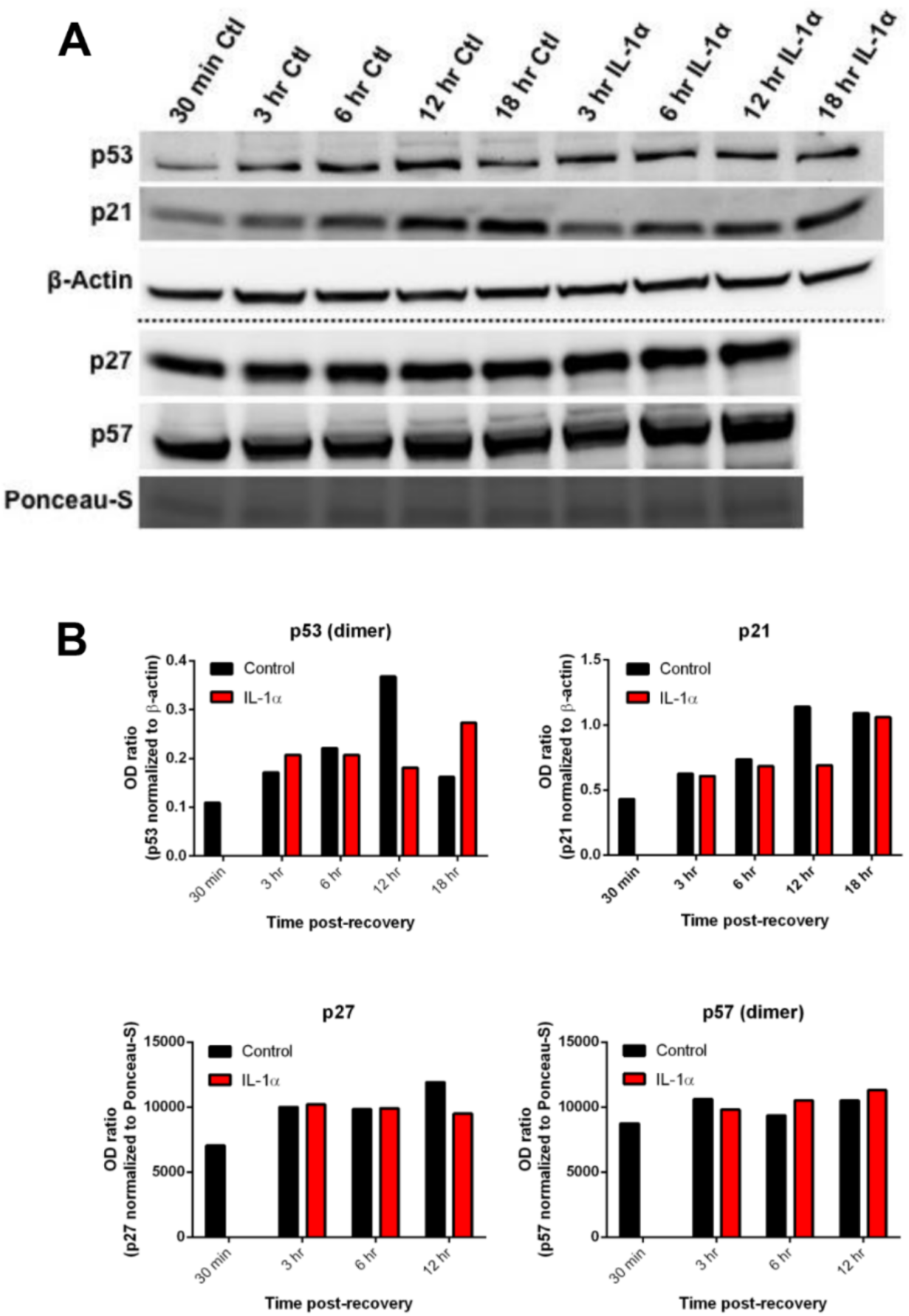
IL-1 does not increase or decrease cell cycle inhibitors in neonatal mouse hippocampal NPs. To begin to elucidate the mechanisms by which IL-1 may be affecting the cycling of neonatal mouse hippocampal NPs, synchronized primary neurospheres were treated with 500 pg/mL IL-1α versus no treatment and cells were harvested for Western blot analysis after 0.5, 3, 6, 12, and 18 hrs and evaluated cell cycle inhibitor levels (Figure S4A). The bands for p53 and p21 were normalized to β-actin, while the bands for p27 and p57 were normalized to Ponceau-S. This experiment failed to show either an increase or a decrease in the levels of the cell cycle inhibitors p21, p27, p53, or p57 with IL-1α treatment (Figure S4B).

**Supplementary Table 1:**
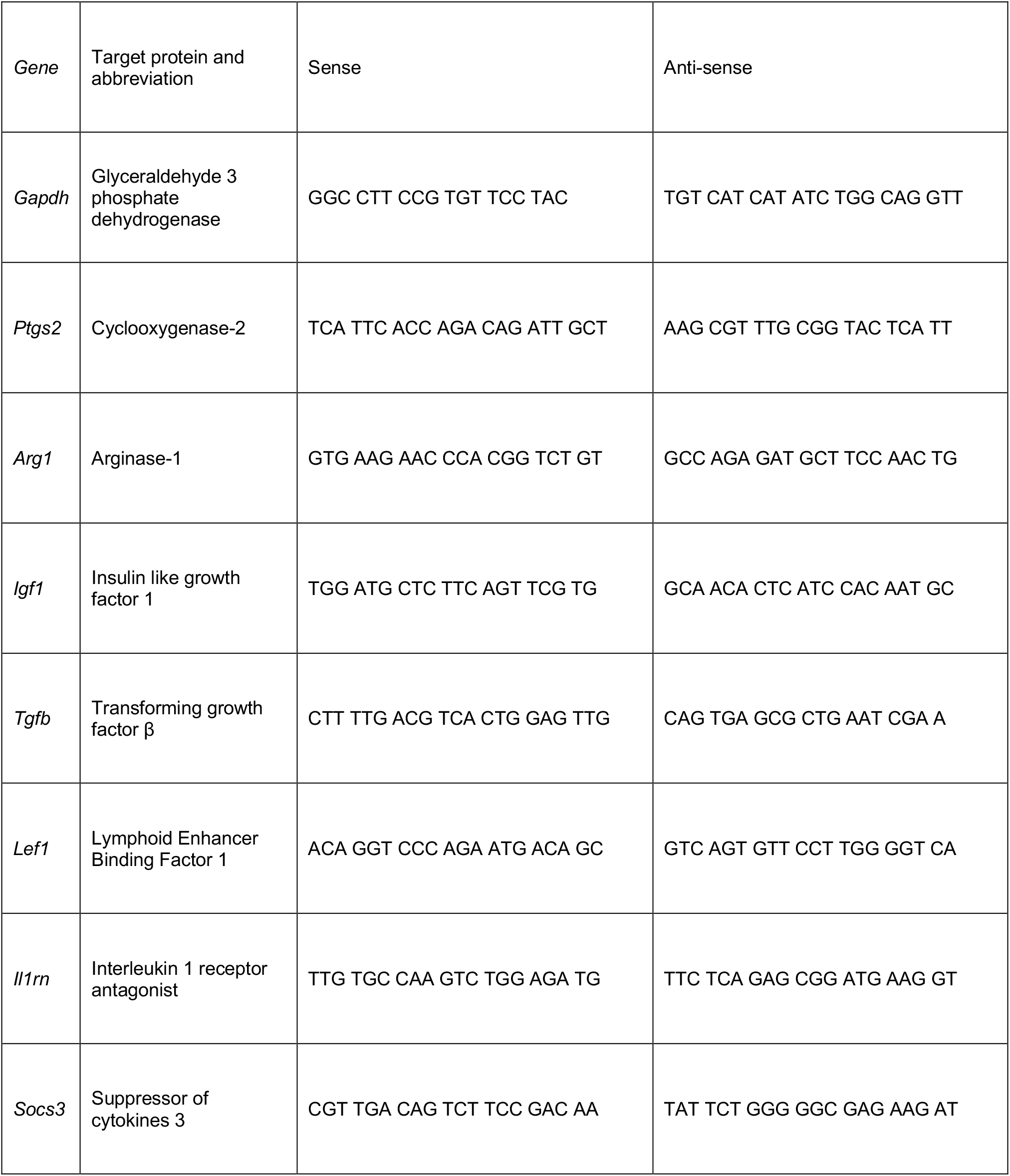
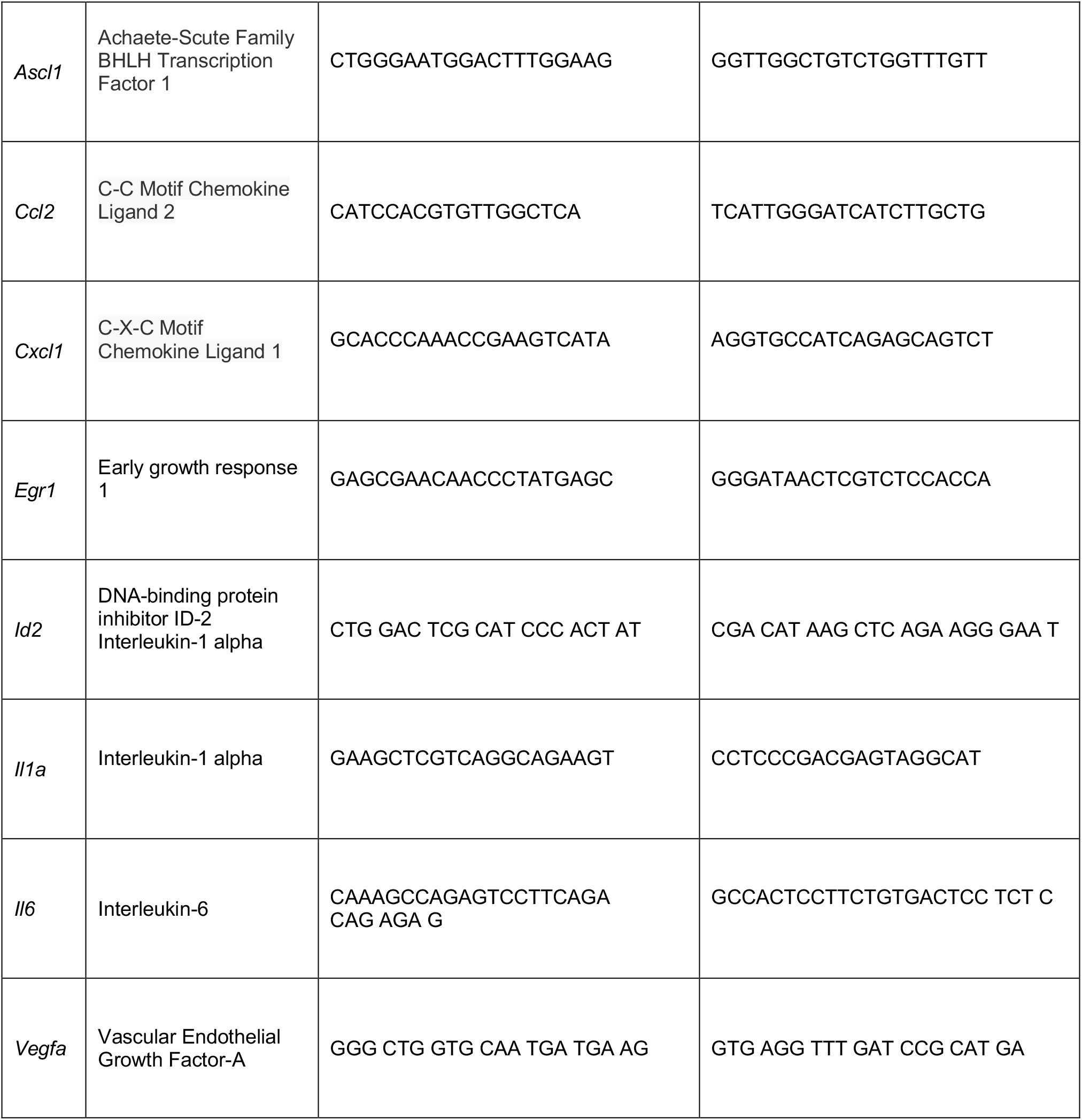
Protein targets and Primer Sequences for Figure S1

